# TPPP Forms Liquid Condensates and Aggregates in Multiple System Atrophy

**DOI:** 10.1101/2023.05.31.542915

**Authors:** Shahrnaz Kemal, Hunter S. Richardson, Thomas S. McAlear, Andrii Kopach, Joseph C. Nowacki, Yan Li, Susanne Bechstedt, Meng-meng Fu

**Author notes:** These authors contributed equally to this work (Shahrnaz Kemal, Hunter S. Richardson, Thomas S. McAlear).

## Abstract

Oligodendrocytes have elaborate arbors of microtubules that extend toward axons and spiral around myelin sheaths. Oligodendrocytes rely on satellite organelles called Golgi outposts to nucleate new microtubules at sites far from the cell body. We now show that the Golgi outpost marker TPPP (tubulin polymerization promoting protein) forms liquid condensates that co-partition with tubulin in order to nucleate microtubules. In oligodendrocytes, TPPP forms either dynamic puncta or aberrant microtubule-associated aggregates. In Multiple System Atrophy (MSA), a sequela of histological events initiates with TPPP aggregation in myelin sheaths and terminates in perinuclear TPPP co-aggregation with alpha-synuclein (aSyn). Finally, recombinant TPPP aggregates are toxic to primary oligodendrocytes. Thus, while the liquid condensate property of TPPP facilitates microtubule nucleation, it also predisposes TPPP to aggregate in disease.

---

Oligodendrocytes extend multiple branching processes that contact neuronal axons then ensheath and wrap them in many concentric layers of myelin. In order to achieve this extraordinary cytoarchitecture, oligodendrocytes rely on a network of microtubules for both structure and transport of organelles and mRNAs (*1, 2*). Proximal radial microtubules are highly branched and found along processes that extend toward the axon. Distal lamellar microtubules spiral from outer to inner layers of the myelin sheath; they are found inside cytoplasmic channels and mediate the transport of vesicular cargo (*3, 4*). Both radial and lamellar microtubules contain Golgi outposts, satellite Golgi organelles that associate with TPPP (tubulin polymerization promoting protein) to nucleate or make new microtubules at sites far from the cell body (*5, 6*). Absence of TPPP results in branching defects, mixed rather than uniform microtubule polarity, and thinner and shorter myelin sheaths. *Tppp* knockout mice have mild motor coordination defects and both innate and learned fear deficits (*5, 7*). Thus, the nucleation of microtubules from Golgi outposts by TPPP is crucial for myelination.

Here, we follow up on histology studies indicating that TPPP localization changes in the degenerative neurological disease MSA (multiple system atrophy) (*8, 9*). Similar to Parkinson’s disease (PD), MSA is a synucleinopathy, meaning that the protein alpha-synuclein (aSyn) aggregates. Unlike PD, in which aSyn aggregates in neurons, MSA harbors glial cytoplasmic inclusions (GCIs). The cell-specific expression of TPPP in oligodendrocytes by RNA-seq (*10*) supports a link between TPPP and GCI formation in MSA. We now demonstrate using biophysical assays, primary oligodendrocyte culture, and human brain histology that TPPP exists as a liquid condensate, a property that is crucial to its function in microtubule nucleation, but predisposes it to aggregation. In primary oligodendrocytes, TPPP also forms liquid condensates that quickly recover after photobleaching, but, when overexpressed, TPPP aggregates along microtubules. In human MSA brains, TPPP aggregation along the myelin sheath then in oligodendrocyte cell bodies likely precedes co-aggregation with aSyn. Finally, recombinant TPPP protein forms fibrils that are toxic to primary oligodendrocytes. Together, these experiments at the molecular, cellular, and brain level demonstrate that the liquid condensate properties of TPPP can become unbalanced, leading to aggregation in neurological disease.

## MSA brains contain perinuclear oligodendrocyte TPPP staining and insoluble TPPP protein

First, we sought to robustly confirm previous histology findings of altered TPPP localization in MSA by increasing the number of patient samples and using high-resolution confocal microscopy. Using immunofluorescent staining of human substantia nigra (SN), we observed striking perinuclear TPPP staining in MSA brains (Fig. 1A, Table S1). We categorized these perinuclear staining patterns by their morphology and intensity as Type 1 (speckled/punctate), Type 2 (dim/diffuse), and Type 3 (bright/compact). We quantified significant differences in non-punctate Type 2 and Type 3 TPPP perinuclear staining when comparing MSA brains to age-matched, sex-matched, and race-matched control brains (Fig. 1B).

**Fig. 1.**
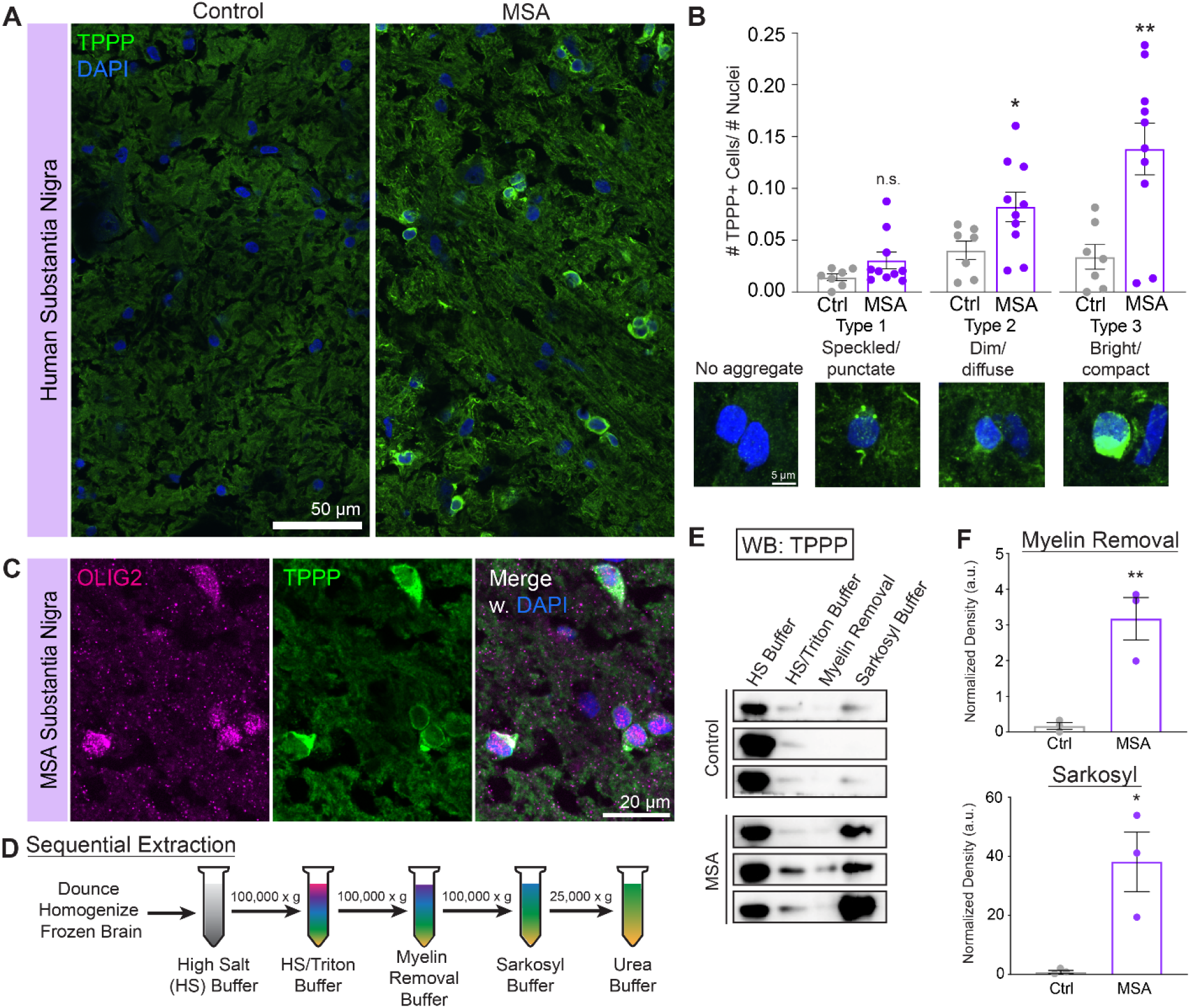
MSA brains contain perinuclear oligodendrocyte TPPP staining and insoluble TPPP protein. (A) Representative micrographs of TPPP staining in SN from control and MSA brains (Patient IDs: S12016, S16219). (B) Density of cells positive for TPPP perinuclear staining normalized by the number of total DAPI-positive nuclei and binned into 3 types of TPPP staining patterns. n = 7, 10 patients; 3–4 images per subject. Type 1 showed no statistical significance (P = 0.126). (C) TPPP staining colocalizes with OLIG2-positive oligodendrocytes (Patient ID: S16023). (D) Schematic of sequential extraction protocol. (E) Western blots of sequential extraction in increasingly stringent buffers of pons from control and MSA brains, immunoblotted against TPPP. n = 3 controls (Patient IDs: S01733, S06589, S08610), 3 MSA (Patient IDs: S11347, S16023, S16219). (F) Quantification of TPPP Western blot bands in myelin-removal and sarkosyl-insoluble fractions. Quantification of HS/Triton fractions showed no statistical significance (P = 0.72). All data represented as means ± SEMs with individual data points. Statistics performed using Student’s *t*-test. *P < 0.05, **P < 0.01, ***P < 0.001.

We further confirmed that these perinuclear TPPP deposits are in oligodendrocytes and colocalize with the marker OLIG2 (*11*) (Fig. 1C). In addition, perinuclear TPPP staining does not colocalize with cell body markers for astrocytes or neurons. We observed no colocalization with the astrocyte marker GFAP (Fig. S1A), the pan-neuronal marker NeuN (Fig. S1B) nor with tyrosine hydroxylase (TH), a marker for dopaminergic neurons, which are abundant in the SN (Fig. S1C).

In order to confirm that TPPP aggregates in MSA, we performed sequential extraction experiments with fresh-frozen tissue from the pons, which is another affected region in MSA. We used increasingly stringent buffers in the following order: high-salt buffer (HS), HS with 1% TX-100, myelin removal buffer, sarkosyl buffer, and urea buffer (Fig. 1D) according to published protocols (*12*). By Western blot, we observed a striking increase in intensity of TPPP bands in both the myelin removal fraction and the sarkosyl fraction from MSA brains when compared to control brains (Fig. 1, E and F, S1D, Table S1). These results indicate that aberrantly high levels of TPPP are present in the myelin fraction in MSA brains. Importantly, they also show that even in the presence of the detergent sarkosyl, TPPP aggregates remain intact in MSA brains.

## TPPP liquid condensates nucleate microtubules

Many sarkosyl-insoluble proteins that aggregate in neurological disease, such as TDP-43 (*12–14*) and FUS (*15, 16*) in amyotrophic lateral sclerosis (ALS), form liquid condensates. Proteins that form liquid condensates typically contain an intrinsically disordered region (IDR). Interestingly, TPPP contains a ∼50-amino-acid N-terminal region with high predictions of disorder by multiple bioinformatics tools, including DISOPRED3 (*17*), MFDp2 (*18*), and PrDOS (*19*) (Fig. 2, A and B). In addition, the machine learning program FuzDrop (*20*) predicts that TPPP has a high probability of forming liquid condensates or undergoing liquid-liquid phase separation (LLPS) (Fig. S2, A–C).

**Fig. 2.**
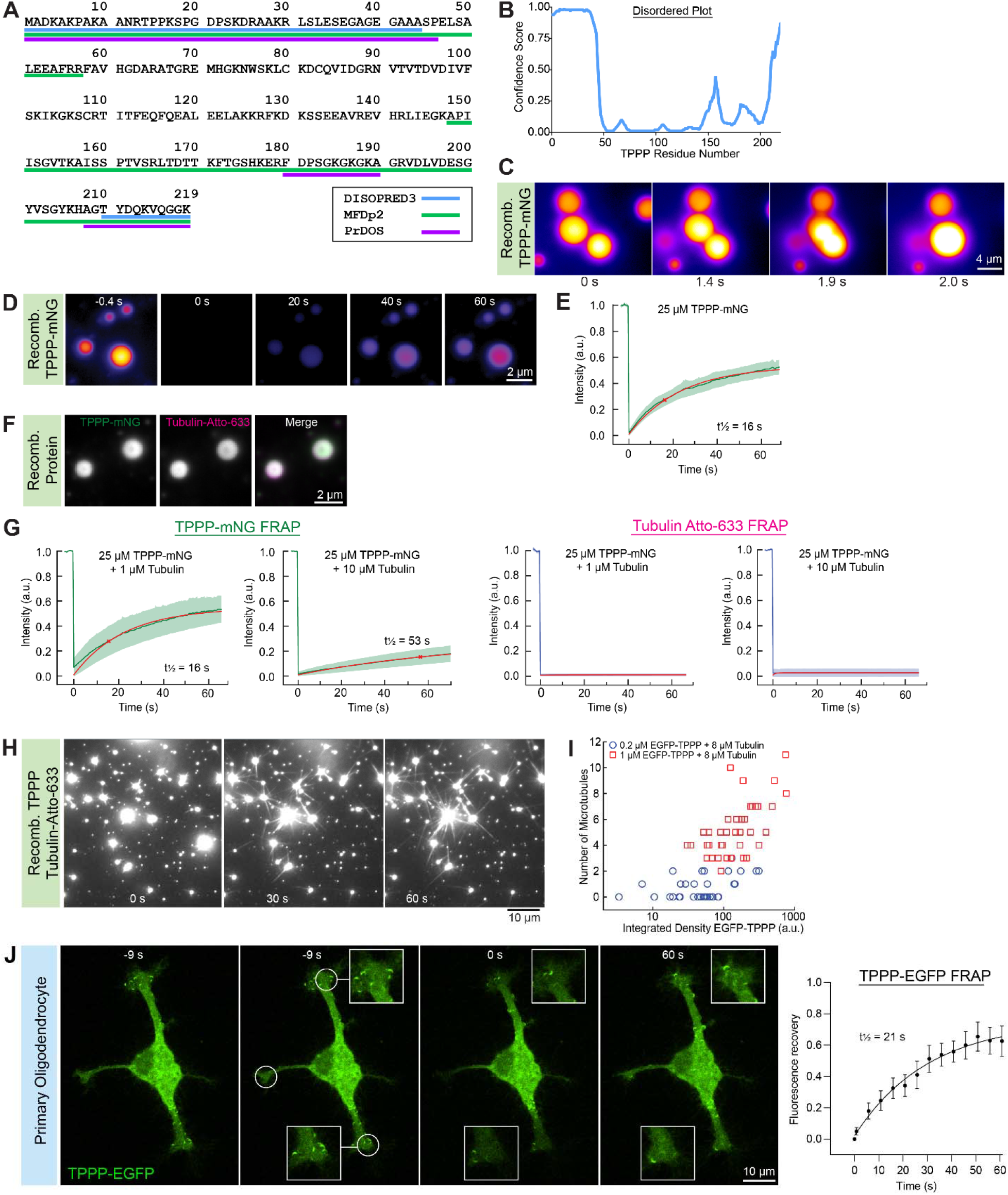
TPPP forms liquid condensates and nucleates microtubules. (A) Bioinformatics predictions of disorder for human TPPP using DISOPRED3, MFDp2, and PrDOS. (B) Confidence score plot of TPPP disorder using DISOPRED3 and residue-based droplet-promoting probability using FuzDrop. (C) Fast fusion of TPPP-mNG liquid droplets occurs on the timescale of ∼1 s. Images acquired at 25 μM TPPP-mNG with 10% PEG 8000 in BRB80. Images in Fig. 2C and 2D are pseudo-colored (yellow = high intensity; purple = low intensity). (D) Representative fluorescent images of 25 µM TPPP-mNG in BRB80 +10% PEG before and after FRAP. (E) Representative FRAP plot of TPPP-mNG signal from Fig. 2D. Time of bleaching is defined as 0 s. n = 28 experiments. Data in Fig. 2E and 2G are represented as mean ± SD (green) and fit to a single exponential y(t)=A(1-e^-t*t^)+c (red). (F) Tubulin partitions into TPPP condensates. 25 µM TPPP-mNG and 2 µM tubulin-Atto633 in BRB80 with 10% PEG 8000 (without GTP). (G) FRAP recovery curves of 25 µM TPPP-mNG in the presence of 1 or 10 µM tubulin (n = 11 FRAP events, 3 experiments). FRAP recovery curves of 1 or 10 µM tubulin-Atto-633 in the presence of 25 µM TPPP (n = 12 FRAP events, 3 experiments). (H) TIRF images of TPPP condensates nucleating microtubules (2 µM unlabeled TPPP, 10 µM tubulin-Atto-633, 10% PEG 8000). See also Movie S1. (I) Scatter plot of EGFP-TPPP integrated density *vs*. number of microtubules nucleated per condensate. Blue circles represent 0.2 µM EGFP-TPPP with 8 µM tubulin condition (n = 35 condensates, 3 experiments). Red squares represent 1 µM EGFP-TPPP with 8 µM tubulin (n = 38 condensates, 3 experiments). (J) FRAP of cultured primary oligodendrocytes expressing punctate TPPP-EGFP for 12–16 hours. White circles indicate photobleached regions. Insets of two regions are presented in white squares. FRAP curve of 12 photobleached regions from 12 cells from 3 separate experiments/animals. Data represented as mean ± SEM. See also Movie S2.

Thus, we asked whether TPPP forms liquid condensates and looked for associated properties of fusion, fluorescence recovery after photobleaching (FRAP), and sensitivity to high salt concentrations. Indeed, we observed that both recombinant TPPP-mNeonGreen (mNG) and EGFP-TPPP (Fig. S2D) form liquid droplets that are up to several μm in diameter (Fig. 2C, S2E). TPPP-mNG droplets fuse rapidly with each other on the timescale of ∼1 s (Fig. 2C). Furthermore, TPPP-mNG droplets recover rapidly after photobleaching with half-time (t_1/2_) of 16 s (Fig. 2, D and E). Interestingly, the N-terminally tagged EGFP-TPPP droplets fuse on a longer time scale (Fig. S2E) and recover less quickly after photobleaching (t_1/2_ = 38s, Fig. S2F), perhaps indicative of the contribution of the N-terminal IDR in liquid condensate dynamicity. Finally, in increasing concentrations of KCl, TPPP condensates shrink in size both in solution and in surface-wetting assays at the imaging surface (Fig. S2G).

As a microtubule-associated protein (MAP), TPPP interacts with tubulin and, thus, we performed experiments to test the association of these proteins in the absence of GTP. In the condition lacking TPPP, free tubulin is soluble. With the addition of TPPP, we observe co-partitioning of TPPP and tubulin (Fig. 2F), indicating the formation of heterogeneous liquid condensates. At a high concentration of TPPP (25 μM) and low concentration of tubulin (1 μM), TPPP rapidly recovers (t_1/2_ = 16 s), which is very similar to its half-time in the absence of tubulin (t_1/2_ = 16 s). However, at a higher concentration of tubulin (10 μM), TPPP recovery decreases (t_1/2_ = 53 s). Surprisingly, in both conditions, tubulin does not recover (Fig. 2G); this observation was very perplexing to us and perhaps indicates the formation of stable structures, such as nascent microtubules, inside these heterogeneous droplets even in the absence of GTP.

Next, we performed microtubule nucleation assays. Previously, we tested microtubule nucleation using bead-associated TPPP to mimic the membrane and geometry of Golgi outposts (*5*). Thus, to test how microtubule formation may occur independent of beads, we reconstituted microtubule dynamics in an *in vitro* total internal reflection fluorescence (TIRF) microscopy assay. In the presence of GTP (1 mM) and low tubulin (10 μM), at which microtubules typically do not form in the presence of tubulin alone, we observed rapid template-independent nucleation of microtubules from TPPP assemblies on the timescale of seconds (Fig. 2H, Movie S1). We also performed similar assays in the presence of recombinant EB3-GFP, which localizes to the outside of the microtubules in the nucleating asters (Fig. S2H). These nucleating asters are symmetric with microtubules emanating in all directions, similar to reconstituted centrosomes (*21*), but different from the branching nucleator augmin, which forms fan-shaped structures (*22*).

We further tested the ability of TPPP to nucleate microtubules at varying concentrations of TPPP and tubulin. First, we lowered the concentration of both TPPP and tubulin and removed PEG. Strikingly, we observed that at concentrations as low as 0.2 μM TPPP and 8 μM tubulin, droplets formed that were able to nucleate new microtubules (Fig. S2I). This concentration is similar to the minimal concentration needed to nucleate microtubules from the γ-tubulin ring complex (γ-TuRC) *in vitro* (*23*). When we increased TPPP to 1 μM, we observed even greater microtubule nucleation (Fig. S2I). Thus, we quantified the integrated density of TPPP and tubulin droplets and we observed that higher TPPP concentrations produced larger TPPP/tubulin droplets (Fig. S2J), which partitioned more tubulin (Fig. S2K), and nucleated more microtubules (Fig. 2I; S2, L and M). Taken together, these experiments suggest that the liquid condensate properties of TPPP are critical for microtubule nucleation.

## Oligodendrocytes contain both TPPP liquid condensates and aggregates

To investigate the liquid condensate properties of TPPP in cells, we turned to primary oligodendrocytes purified from rat brains using the immunopanning technique (*24*). We expressed TPPP-EGFP in oligodendrocytes and observed small, motile EGFP-positive puncta throughout the cytoplasm (Movie S2). We found that expression for longer than 24 hours was detrimental to overall health and viability of the oligodendrocytes (Fig S3A). Therefore, we confined our analysis to 12–16 hours post-transfection. These TPPP-EGFP puncta exhibited fast recovery after photobleaching on the timescale of <1 min and t_1/2_ = 21 s (Fig. 2J, Movie S2).

Surprisingly, higher expression levels of TPPP-EGFP were associated with aberrant localization in a filamentous pattern reminiscent of microtubules (Fig. 3A). Colocalization of TPPP-EGFP with microtubules was confirmed by fixation and immunostaining for α-tubulin (Fig. S3A). These images and line scan analyses show that filamentous TPPP does not perfectly colocalize with microtubules, but instead appear to accumulate on top of and around microtubules, indicating that they are not uniformly integrated into or onto the microtubule lattice. Remarkably, TPPP-EGFP found in this filamentous pattern did not recover in FRAP experiments (Fig. 3A, Movie S3), even when imaged for up to 10–15 min. Together with our *in vitro* FRAP experiments of co-partitioning TPPP and tubulin droplets, which did not exhibit tubulin recovery (Fig. 2E), this suggests that association with tubulin decreases TPPP dynamicity and increases its propensity to aggregate.

**Fig. 3.**
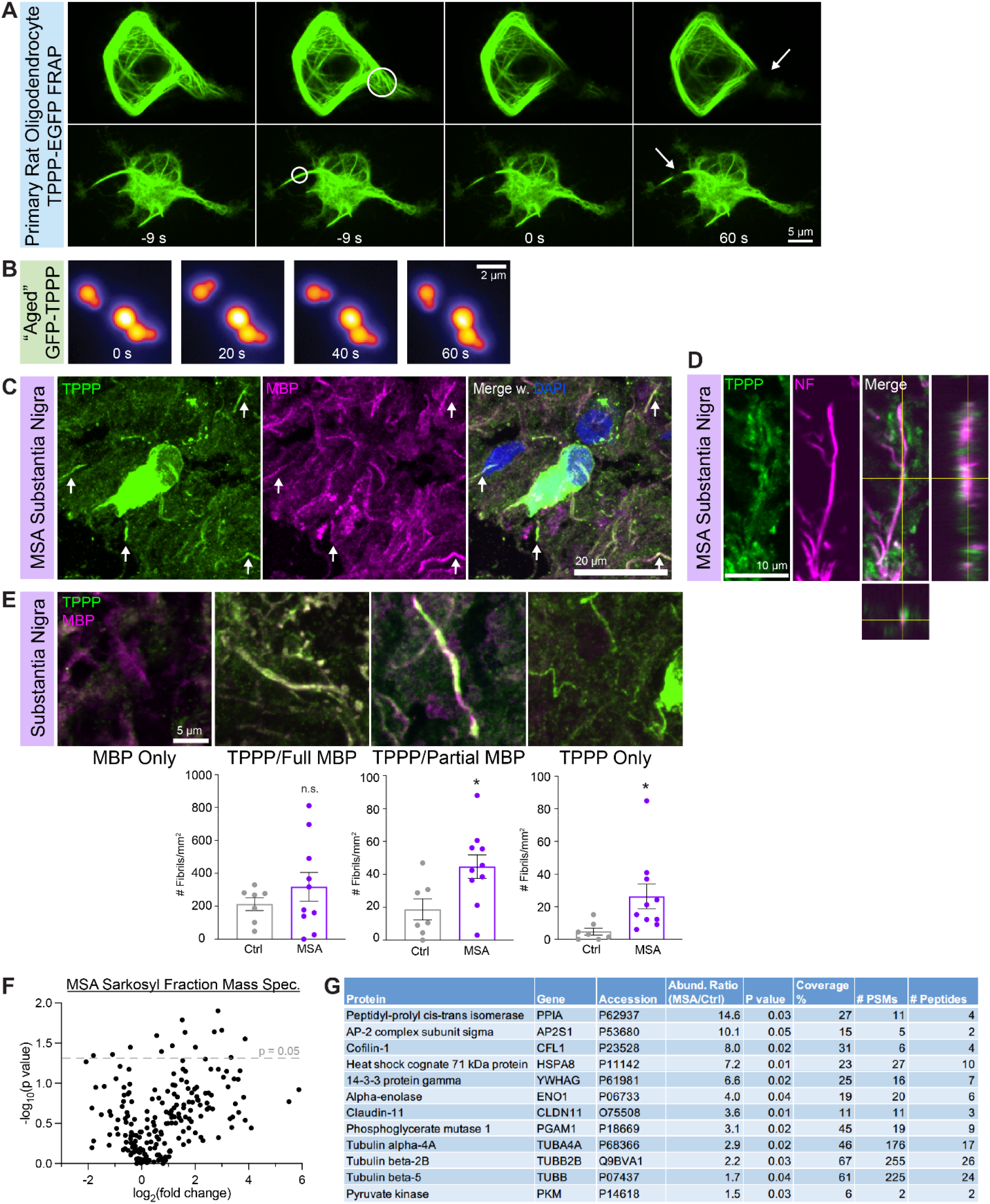
TPPP fibrils form in the myelin sheath. (A) FRAP of cultured primary oligodendrocytes expressing filamentous TPPP-EGFP for 12–16 hours. See also Movie S3. (B) Biophysical “aging” assay with EGFP-TPPP (10 µM) in BRB80 + 100 mM KCl + 10% PEG 8000 incubated at 35°C for 1 hour. These droplets no longer fuse rapidly and form stable “shmoo” structures. (C) Representative image of fibrillar TPPP staining that colocalizes with the myelin sheath marker MBP from MSA SN (Patient ID: S03266). (D) Localization of TPPP fibrils that spiral around the axon marker NF in MSA SN (Patient ID: S11347). (E) Classification of TPPP and MBP colocalization into 4 categories: MBP only, TPPP/full MBP colocalization, TPPP/partial MBP colocalization, and TPPP only. Quantification of TPPP and MBP colocalization by number of fibrils per square millimeter. n = 7 control SN, 10 MSA SN, 1–2 images per subject. Student’s *t*-test, *P < 0.05. Full MBP comparison showed no statistical significance (P = 0.354). (F) Volcano plot comparing fold change of 210 proteins identified from sarkosyl fractions from MSA pons versus control pons. Fold change (normalized abundance ratios) and p-values (ANOVA) were calculated using Proteome Discoverer software. (G) List of proteins significantly enriched in MSA pons sarkosyl fractions compared to control. n = 3 control pons, 3 MSA pons.

Because the filamentous, microtubule-associated TPPP may represent aberrant liquid-solid transitions or a gel-like state, we also turned to biophysical FRAP assays and aging assays using recombinant TPPP to better understand these transitions. Because we previously noted that N-terminally tagged EGFP-TPPP exhibits less efficient FRAP recovery (t_1/2_ = 38 s, Fig. S2F), we added increasing concentration of tubulin. We observed that with increasing concentrations of tubulin, FRAP recovery also decreased (2 μM tubulin: t_1/2_ = 81 s, 10 μM tubulin: t_1/2_ = 203 s, Fig. S3B). This suggests that concentrations of tubulin may play a role in TPPP aggregation in cells. For aging assays, we incubated TPPP at 37°C for 1 hour with no shaking and observed the formation of aberrant non-spherical shapes (Fig. 3B). These shapes resemble yeast shmoos and are likely droplets mid-fusion, but they do not fuse even after >1 min. These results indicate that TPPP is prone to liquid-solid transitions on short timescales and that adjacent liquid condensates that fail to fuse may represent the very early stages of aggregation.

## TPPP fibrils form in the myelin sheath

The aberrant colocalization of aggregate TPPP and microtubules in oligodendrocytes led us to further investigate fibrillar TPPP staining that we observed in MSA brains (Fig. 3C). Because endogenous TPPP puncta localize to Golgi outposts found along microtubules in oligodendrocytes (*5*), we hypothesized that these TPPP fibrils are also found in oligodendrocytes. We first confirmed that these TPPP fibrils are not located inside the processes of astrocytes or neurons. We confirmed by staining that TPPP fibrillar aggregates neither colocalize with the astrocyte marker GFAP (glial fibrillary acid protein) (Fig. S1A) nor with the neuronal marker neurofilament (NF) (Fig. S3C). We wanted to further distinguish between the two types of microtubules in oligodendrocytes: radial microtubules that project toward axons and lamellar microtubules that traverse outer and inner layers of the myelin sheath. Finally, using the myelin sheath marker MBP (myelin basic protein), we found robust colocalization with TPPP (Fig. 3C). Importantly, co-staining shows that TPPP fibrils can spiral around NF-positive axons (Fig. 3D) and morphologically resemble the cytoplasmic channels inside myelin sheaths that contain microtubules and facilitate transport (*3, 4*). These results are consistent with high levels of TPPP protein in the myelin fraction of MSA brains in sequential extraction experiments (Fig. 1, E and F).

Previous histology studies have hypothesized that TPPP aggregation may begin inside the cell body, then spread to the myelin sheath. We tested this by categorizing the colocalization between TPPP and MBP. We binned fibrillar TPPP staining into three categories: fully colocalized with MBP, partially colocalized with MBP, and TPPP only. (Fig. 3E). Surprisingly, we found no difference in density of TPPP fibrils that are fully colocalized with MBP when comparing between age-matched control brains and MSA brains. This perhaps indicates a propensity for TPPP to form fibrillar structures in aging brains, regardless of disease status. However, in MSA brains, we indeed observe significant increases in density of TPPP fibrils that are partially colocalized with MBP or not colocalized with MBP (Fig. 3E), which we interpret to indicate partially or fully degenerated myelin sheaths, respectively. This quantification, together with increases in perinuclear TPPP staining in MSA, indicate that TPPP fibrils along the myelin sheath likely precedes perinuclear accumulation in the cell body.

Finally, we performed mass spectrometry of sarkosyl fractions from MSA brains (Fig. 1D–F) and confirmed that several tubulin proteins (TUBA4A, TUBB2B, TUBB) are indeed enriched in MSA versus control fractions (Fig. 3, F and G). Furthermore, the tight-junction protein claudin 11 that is highly and specifically expressed by oligodendrocytes (*10, 25*) is also abundant in MSA sarkosyl fractions. These findings are consistent with our cell culture and histology results that support aggregation of TPPP in microtubule-rich cytoplasmic channels found along myelin sheaths. In addition, MSA sarkosyl fractions are also enriched in chaperone proteins (PPIA, HSPA8), perhaps indicative of an upregulation response to misfolded and aggregated proteins. Surprisingly, MSA sarkosyl fractions are also enriched in several members of the glycolysis pathway (ENO1, PGAM1, PKM), perhaps indicative of metabolic stress. Together, our proteomics results paint a picture of an aggregation microenvironment undergoing metabolic and protein folding stress.

## Many perinuclear TPPP accumulations do not colocalize with aSyn in MSA

Because glial aSyn aggregates are the histological hallmark and postmortem diagnostic criteria of MSA, we compared perinuclear TPPP staining versus perinuclear aSyn staining. First, we confirmed that there are more aSyn and TPPP perinuclear accumulations in MSA SN than control tissue (Fig. S4, A and B), consistent with our earlier quantifications (Fig. 1B). In merged images, we observe three populations of perinuclear staining: TPPP only, colocalized TPPP/aSyn, and aSyn only (Fig. 4A). We quantified significantly more TPPP/aSyn and aSyn-only perinuclear staining in both MSA SN compared to control tissues (Fig. S4C), because control SN has very few to no aSyn perinuclear staining (Fig. S4, D and E). Surprisingly, we found that MSA SN contains significantly more TPPP-only accumulations than TPPP/aSyn or aSyn-only accumulations (Fig. 4B), when normalized by number per nuclei. Looking at the distribution of these three populations in MSA, ∼55% are TPPP-only, ∼35% are TPPP/aSyn colocalized, and only ∼10% are aSyn-only (Fig. 4C). Taken together, these data suggest that the TPPP only subpopulation may present an earlier stage of disease.

**Fig. 4.**
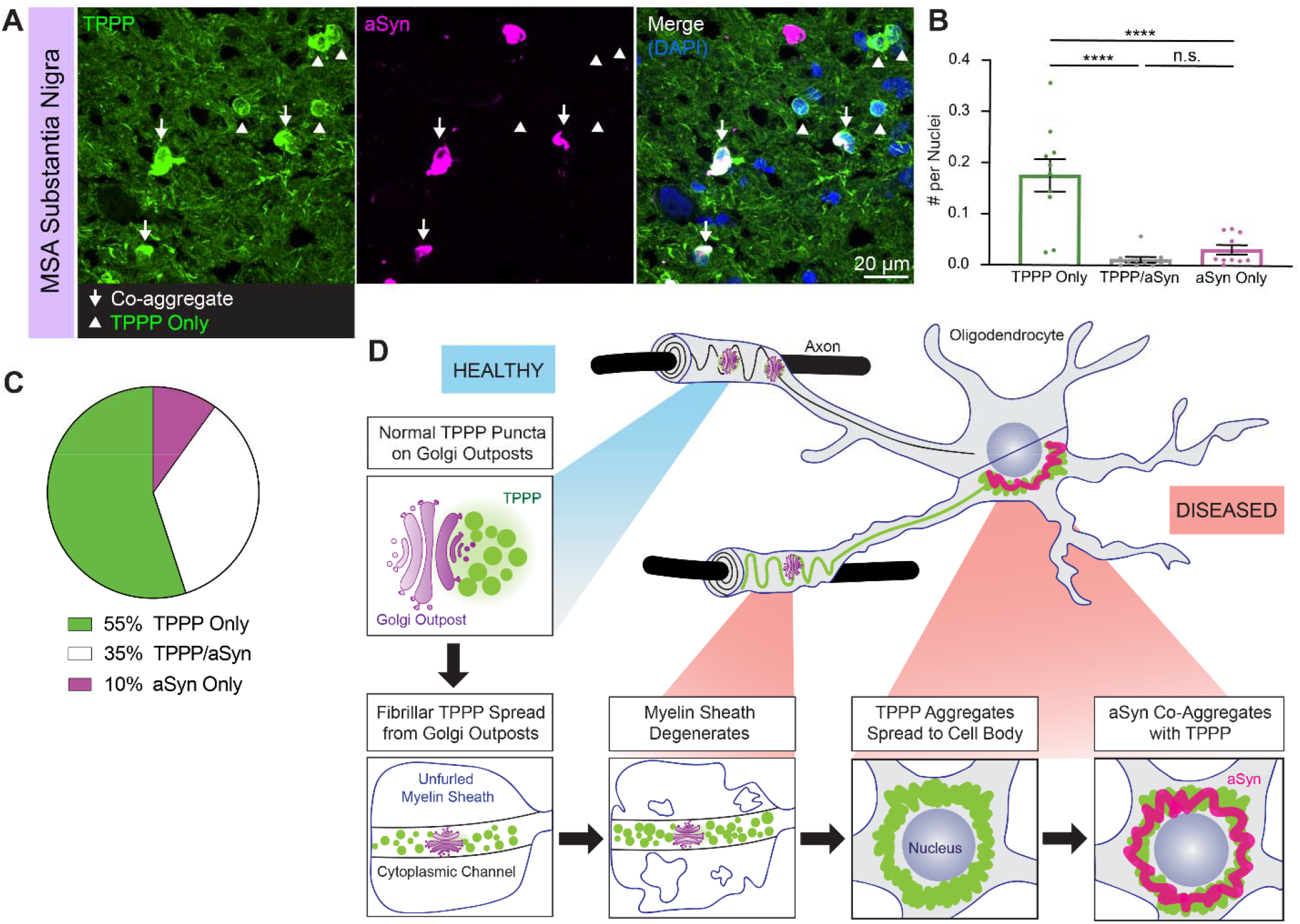
Many perinuclear TPPP accumulations do not colocalize with aSyn in MSA. (A) Representative image of perinuclear TPPP and aSyn aggregates in MSA SN (Patient ID: S16219). (B) Quantification of TPPP and aSyn colocalization by number. n = 10 MSA SN; 2 images per subject. One-way ANOVA: ****P < 0.0001; n.s. (P = 0.741). (C) Pie chart illustrating percentages/distribution of all TPPP/aSyn perinuclear staining in MSA SN. (D) Schematic showing proposed pathological cascade beginning with normal TPPP punctate localization on Golgi outposts. With age, fibrillar TPPP forms in the myelin sheath, but, in MSA, the myelin sheath degenerates. Later, perinuclear TPPP staining is observed, then aSyn co-aggregation.

Together, our histological, cellular, and biophysical data suggest a cascade of pathological events in MSA (Fig. 4D). Initially, TPPP liquid condensates associated with Golgi outposts have punctate expression along myelin sheaths. With age, fibrillar TPPP accumulates along microtubules in the myelin sheath. In MSA patients, myelin sheaths containing fibrillar TPPP begin to degenerate and aggregate TPPP spreads toward the cell body. As perinuclear TPPP aggregates accumulate, they then begin to co-aggregate with aSyn, which has been the pathological diagnostic criteria of MSA for decades.

## aSyn is expressed by oligodendrocytes and co-partitions in liquid condensates

Previous mechanistic studies of aSyn function indicate that neuronal aSyn participates in synaptic exocytosis (*26, 27*). Thus, we wondered whether the source of aSyn that colocalizes with perinuclear TPPP in oligodendrocytes is from neuronal release or endogenous oligodendrocyte production. Indeed, staining in cultured oligodendrocytes (*28*) indicates that oligodendrocytes themselves also express aSyn. Thus, we validate by different methods that aSyn is expressed by oligodendrocytes. First, a bulk RNA-seq database of neonatal rodent brain cells acutely isolated using the immunopanning technique (*10*) indicates abundant expression of *aSyn* mRNA by oligodendrocyte precursor cells (OPCs) and oligodendrocytes (Fig. S5A). An additional bulk RNA-seq database of immunopanned brain cells from human brains (*29*) also indicates high levels of *aSyn* mRNA expression by oligodendrocytes (Fig. 5A). Second, western blotting of differentiated primary rat oligodendrocyte cell lysate shows the presence of aSyn protein at levels comparable to that in cultured primary hippocampal neurons (Fig. 5B). Third, staining of primary immunopanned rat oligodendrocytes shows punctate aSyn throughout the cell (Fig. 5C). Comparing across different developmental time points of differentiation, aSyn signal is most robust in pre-myelinating oligodendrocytes prior to expression of MBP protein and as MBP protein translation begins (Fig. S5B). Fourth, aSyn staining is also present in primary rodent oligodendrocytes cultured on 3D microfibers as they begin to make nascent myelin sheaths (Fig. 5D).

**Fig. 5.**
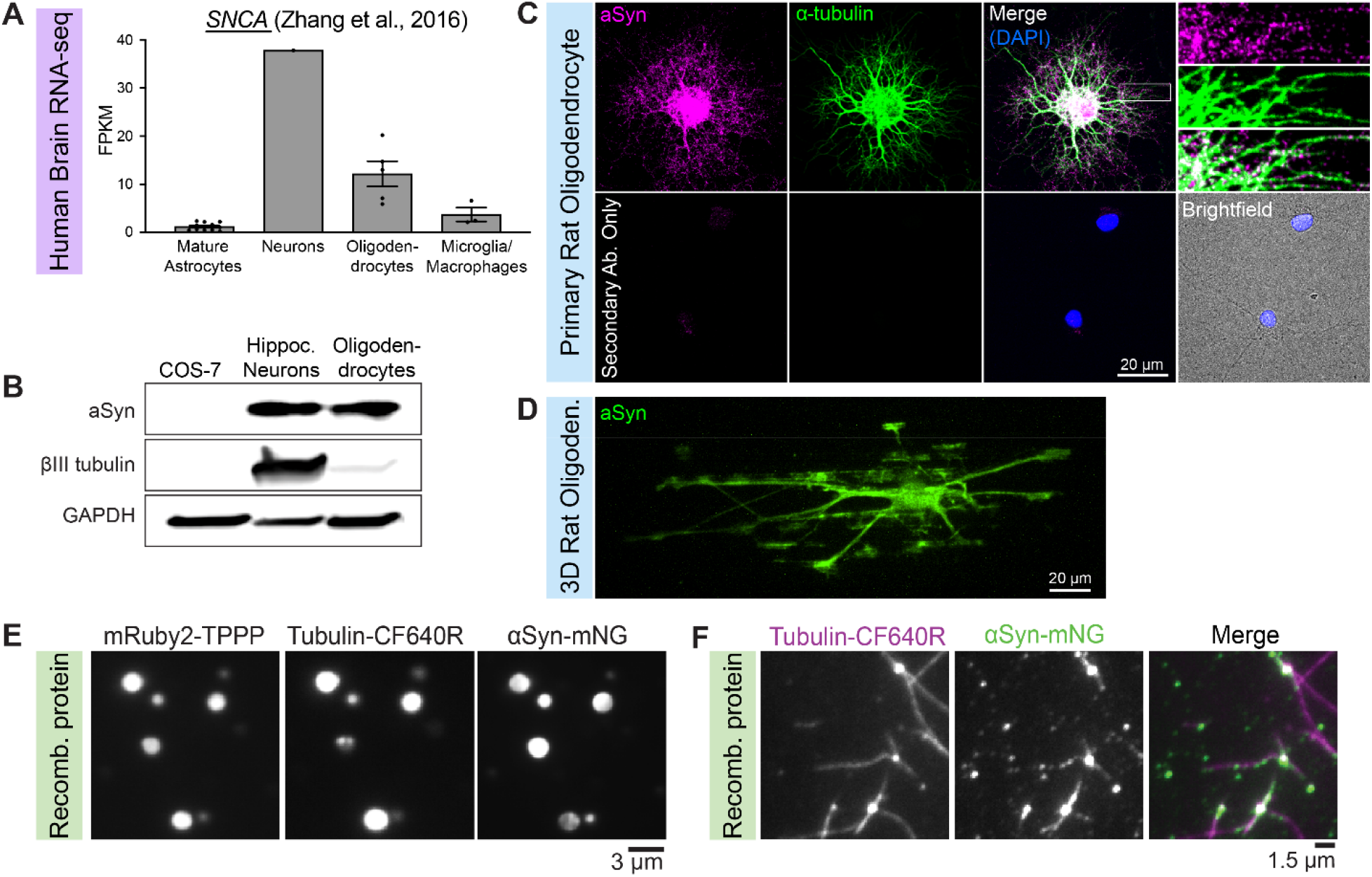
aSyn is Expressed by oligodendrocytes and co-partitions in liquid condensates. (A) *SNCA* (aSyn) mRNA expression in a bulk RNA-seq database of immunopanned brain cells from *Homo sapiens* brains (*29*). FPKM (fragments per kilobase of transcript per million mapped reads). (B) Primary rat oligodendrocyte differentiated for 6 days *in vitro* stained against aSyn and tubulin. The secondary-antibody-only control shows no staining in regions containing cells whose contours can be visualized in brightfield. (C) Western blot comparing aSyn protein levels in lysates from COS-7 cells, primary rat hippocampal neurons, and primary rat oligodendrocytes. This βIII tubulin antibody labels a tubulin that is enriched in neurons but not in oligodendrocytes. (D) Immunostaining against aSyn in primary rat oligodendrocyte cultured on biologically inert 3D microfibers that mimic the shape of axons. (E) Fluorescent images of TPPP, aSyn, and tubulin co-condensates. 3.5 µM mRuby2-TPPP, 2 µM aSyn-mNG, 2 µM tubulin-CF640R in BRB80 with 10% PEG 8000. (F) TIRF microscopy images of TPPP condensates nucleating microtubules in the presence of aSyn. 1 µM TPPP, 0.2 µM aSyn-mNG, 8 µM tubulin-CF640R in BRB80.

To better understand how aSyn may interact with TPPP, we turned to experiments both in primary oligodendrocytes as well as biophysical assays with recombinant protein. First, we co-expressed TPPP-EGFP and aSyn-mRuby in primary oligodendrocytes and observed colocalization of TPPP-EGFP and aSyn-mRuby (Fig. S5C). Second, we asked how aSyn may affect the liquid condensate properties of TPPP and tubulin using recombinant proteins in biophysical assays. Indeed, aSyn forms liquid condensates at high concentrations ∼200 μM (*30*), while TPPP forms liquid condensates at a much lower μM range. We first asked whether aSyn and tubulin co-partition in the absence of TPPP. We found that tubulin remained soluble at two different concentrations of aSyn (Fig. S5D). Next, we added TPPP and found that all three proteins now co-partitioned together, forming heterogeneous liquid condensates (Fig. 5E). Finally, we asked how these proteins might interact with each other in microtubule nucleation assays. In the presence of GTP, we found that TPPP continued to nucleate microtubules. Surprisingly, though, we observed some aSyn decoration of microtubules in a fibrillar pattern proximal to the liquid condensate and in a punctate pattern more distally (Fig. 5F). Together, these results indicate that aSyn and TPPP are located in close proximity in oligodendrocytes and that they interact *in vitro*. These data indicate that TPPP may recruit aSyn, bringing together two aggregation-prone proteins at the molecular level.

## TPPP pre-formed fibrils (PFFs) are toxic to oligodendrocytes

Though our data so far strongly support the correlation between TPPP aggregation and disease, we next tested the causation between TPPP aggregation and disease. To determine whether TPPP has an intrinsic propensity to aggregate, we aimed to reconstitute this process in a test tube. First, we quantified endogenous levels of TPPP in rodent brain lysate and primary oligodendrocytes using recombinant TPPP as a standard. We estimate TPPP concentrations ∼11 nM in the brain and ∼29 nM in oligodendrocytes (Fig. S6A), but that local concentrations at Golgi outposts are likely in the μM range. Next, we generated PFFs by shaking 7 μM unlabeled recombinant TPPP (Fig. S6B) for 24–48 hours at 37°C in PBS (phosphate buffered saline). We determined that absorbance of TPPP decreased over time, indicating a decrease in soluble TPPP (Fig. S6C). Importantly, we used much lower concentrations of TPPP and shorter time periods than previous studies that generated aSyn PFFs, which used ∼100–350 μM aSyn for usually 5–7 days (Fig. S6D) (*31–34*).

Next, we used transmission electron microscopy (TEM) to visualize these samples. We detected formation of fibrous assemblies in the reaction volume. Surprisingly, we also observed a second population of denser aggregates (Fig. 6A). We categorized these assemblies as “fibrils” and “chunks”, respectively, based on their morphology. We confirmed using immunogold labeling that both fibrils and chunks are positive for TPPP (Fig. 6A). Interestingly, chunks have a higher density of TPPP-positive gold bead labeling (Fig. 6B), perhaps indicating that the conformation of chunks is different from that of fibrils. Importantly, these assemblies formed fast and at low concentrations of TPPP that are more than one order of magnitude lower than concentrations used for formation of aSyn PFFs (Fig. S6D). Taken together, these data indicate that TPPP is extremely prone to aggregation.

**Fig. 6.**
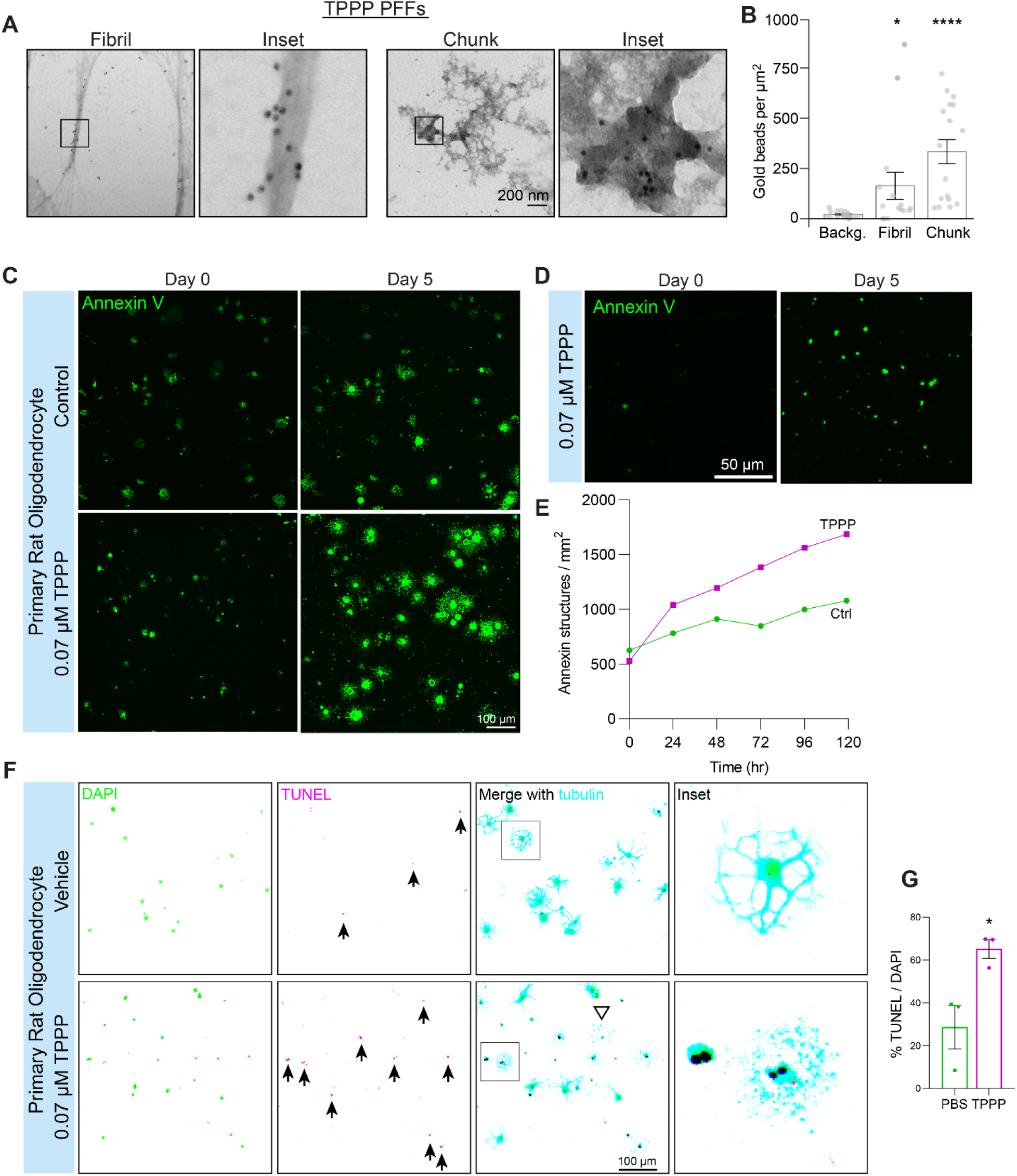
TPPP pre-formed fibrils (PFFs) are toxic to oligodendrocytes. (A) Representative TEM micrographs of immunogold labeled fibrous or chunky assemblies composed of TPPP. Magnified insets show that gold beads colocalize with these TPPP assemblies. (B) Quantification of immunogold labeling normalized by area. N = 20 (background control images), 15 (fibrils), 17 (chunks). Kruskal-Wallis test; *P < 0.05, ****P < 0.0001. (C) Long-term live-cell imaging of primary rat oligodendrocytes treated with 0.07 µM TPPP PFFs and annexin V dye. (D) Annexin-positive blebs in TPPP PFF-treated oligodendrocyte cultures. (E) Quantification of annexin-positive structures. (F) TUNEL staining of differentiated primary rat oligodendrocytes treated with 0.07 µM TPPP PFFs. Black arrowheads indicate TUNEL-positive cells. White triangles indicate DAPI-negative TUNEL-negative dead “shadow” cells in which the nuclei have detached. (G) Quantification of TUNEL-positive nuclei as a percent of all DAPI positive nuclei. n = 3 experiments, 627 nuclei (PBS-treated), 995 nuclei (TPPP-treated). Student’s *t*-test: *P < 0.05. All data represent mean ± SEM.

Finally, we asked if TPPP PFFs can induce cell death in primary rat oligodendrocytes. We sonicated TPPP PFFs to produce smaller PFFs with diameter ∼67-nm (Fig. S6E and F). We performed long-term live-cell confocal imaging experiments in which primary oligodendrocytes were incubated with 0.07 μM TPPP PFFs. We did not add any chemical or liposomal transfection reagents. We used a fluorescent annexin V dye to detect phosphatidylserines that allow visualization of cell death. Though we see some annexin-positive cells in the control condition too, this is expected based on programmed cell death in oligodendrocytes (*35*). In the TPPP PFF condition, we observed rapid conversion to annexin-positive cells as early as one day of incubation (Fig. 6, C and E, Movie S4). In addition, we observed the dispersion of small annexin-positive puncta (Fig. 6D), which are likely blebs from dying cells (*36*). We further confirmed cell death using TUNEL staining of differentiated primary oligodendrocytes incubated with different concentrations of TPPP PFF (Fig. 6, F and G; Fig. S6F). The lower concentrations of TPPP (0.07 µM and 0.007 µM) are closer to quantified endogenous levels (Fig. S6A). All concentrations induce TUNEL positivity and result in the appearance of dead and damaged cells. Taken together, these studies indicate that TPPP PFFs can potently kill oligodendrocytes at physiological concentrations.

## DISCUSSION

Here, we show a crucial link between the normal biophysical properties of TPPP and its aberrant aggregation in neurological disease. Similar to other microtubule nucleators in centrosomes (*21*) and such as TPX2 (*37*), recombinant TPPP behaves as a liquid condensate that is dynamic in FRAP assays and partitions tubulin in order to nucleate microtubules. As a Golgi outpost protein, TPPP functions in oligodendrocytes to nucleate microtubules at sites far from the nucleus. In primary oligodendrocytes, punctate TPPP is also dynamic in FRAP assays, but cells over expressing TPPP aberrantly accumulate TPPP along the microtubule lattice. This is consistent with fibrillar populations of TPPP aggregates found along degenerating myelin sheaths in MSA and with PFFs and chunks formed from recombinant TPPP in *in vitro* aggregation assays. Finally, sonicated TPPP PFFs are toxic to primary oligodendrocytes, demonstrating a causative link between aggregate TPPP and cell death.

Possible molecular events that lead to the switch between normal TPPP localization and TPPP aggregation with age include changes in post-translational modifications (PTMs), interactions with other MAPs, and translational control. Other liquid condensate proteins that aggregate in neurological diseases exhibit altered propensity to aggregate based on mutations and post-translational modifications (PTMs) (*38*). Though no known *TPPP* mutations have been characterized, we are actively pursuing investigations of TPPP PTMs. In addition, we previously postulated that TPPP likely functions with partner MAPs (*5*) and ongoing studies in our lab are investigating both the underlying cell biology mechanisms of these interactions as well as their consequences in disease. In addition, classic studies indicate by Western blot that TPPP protein levels increase with age in rats, with highest expression at 1 or 2 years of age (*39*). Modern mass spectrometry studies indicate that TPPP is the 31st most abundant protein in myelin (*25*) and that its expression increases in OPCs isolated from aging rats (*40*). However, this could either be interpreted as a higher demand for TPPP function or a defect in TPPP clearance or possibly a combination of both.

Our histological data suggest a cascade of pathological events beginning with TPPP fibrillar aggregation, which spreads to the cell body, then progresses to co-aggregate with aSyn. Surprisingly, even age-matched control brains contain TPPP-positive MBP-positive fibrillar aggregates, but these are contained within intact myelin sheaths. In MSA brains, TPPP aggregates colocalize with degenerating myelin sheath and strikingly accumulate in the cell body. However, not all perinuclear TPPP aggregates colocalize with aSyn, indicating that aSyn aggregation may represent a late stage of MSA. In other words, TPPP may be an indicator of early disease and therefore is a promising candidate as a biomarker for MSA. Our previous work in *Tppp* knockout mice indicate that complete loss of TPPP leads to fear deficits and mild motor coordination defects, but no observed differences in lifespan (*5, 7*). This may indicate that *TPPP* mRNA is a viable target for knockdown therapies, such as anti-sense oligonucleotides (ASOs) therapies. However, animal studies should first confirm the causation between TPPP aggregation and disease and ongoing experiments in our lab address whether stereotaxically injected TPPP PFFs can lead to pathology and behavior deficits in mice.

Broadly, our study may also have implications on the understanding of other neurological diseases. In other synucleinopathies, such as PD and Lewy body dementia (LBD), TPPP may play a role in disease pathology. In Alzheimer’s disease (AD), recent snRNA-seq analyses indicate significant transcriptional changes in oligodendrocytes (*41*). Indeed, an early histology paper suggests that TPPP may have aberrant fibrillar localization in AD (*42*). In addition, in a form of leukodystrophy, a childhood white matter disease, the tubulin gene *TUBB4A* is mutated (*43*), but it is unclear whether TPPP may be involved in disease pathology. Thus, future studies on the mechanism of TPPP aggregation and efforts to validate TPPP as a biomarker for diverse diseases are timely and necessary.

## METHODS

### Human brain immunohistochemistry

Fresh frozen human brain substantia nigra (SN) with post-mortem intervals <24 h from 10 MSA and 7 control cases were obtained from the Harvard Brain Bank through the NIH NeuroBioBank. Frozen human brain tissue was mounted in Tissue Plus O.C.T. Compound (Fisher HealthCare, 23-730-571) and cryosectioned to 12-µm thick slices.

Cryosections were fixed in 4% paraformaldehyde (Electron Microscopy Services, 15711) for 10 min at room temperature, and blocked with 10% donkey serum in 0.1% Triton X-100 in PBS for 30 min. Cryosections then were incubated at 4°C overnight with the following primary antibodies at these concentrations: aSyn (Abcam ab209420, 1:1000), phosphorylated aSyn (Wako 015-25191, 1:1000), GFAP (Cell Signaling 123895, 1:400), MBP (Abcam ab7349, 1:100), NF (Biolegend 801701, 1:250), NeuN (Millipore ABN78, 1:250), OLIG2 (IBL 18953, 1:250), TPPP (ThermoFisher PA5-19243, 1:250), and/or tyrosine hydroxylase (Abcam ab76442, 1:500). After, the sections were incubated in secondary antibodies (Jackson ImmunoResearch Laboratories, 1:100 or 1:200) for 1 hour at room temperature. A DAPI stain (Sigma D9542) was applied for 3 min before the antibodies were fixed with a 10 min 4% PFA incubation at room temperature. Next, the slides were immersed in 0.2–0.5% Sudan Black B (SBB) (Sigma 199664) for 20 min at room temperature. After washing in 0.02% Tween, the sections were mounted in Vectashield Plus Mounting Medium (Vector Laboratories, H-1900) or Prolong Gold (Invitrogen P36934).

Immunofluorescent images of the substantia nigra pars compacta were acquired on a Nikon Ti2 spinning-disk confocal with Hamamatsu ORCA FusionBT sCMOS camera using 10X and 60X objectives. 1–2 574×574 µm images and 4–6 125×125 µm images with 7.5-10 µm z-stacks were collected for each stain. DAPI spot counting was performed on the larger images using the pixel and object classification functions of the machine learning software ilastik (*44*). Aggregate counting was performed using the Cell Counting plugin from Fiji (*45*).

### Human brain sequential extraction

Fresh frozen human pons from 3 control (S01733, S06589, S08610) and 3 MSA (S11347, S16023, S16219) cases were Dounce homogenized in High-Salt Buffer (750mM NaCl, 50mM Tris-HCl, 10mM NaF, 5mM EDTA), then spun at 100,000 x g for 30 min at 4°C in an ultracentrifuge (Beckman OPTIMA TLX with TLA-100.3 rotor). The supernatant was saved and the pellet went through a series of similar extractions using High-Salt Triton Buffer (High-Salt Buffer, 1% Triton X-100), Myelin Removal Buffer (High-Salt Triton Buffer, 30% sucrose), High-Salt Sarkosyl Buffer (High-Salt Buffer, 1% sarkosyl), and Urea Buffer (7M urea, 2M thiourea, 4% CHAPS, 30mM Tris-HCl). Each step included a 30 min spin at 100,000 x g and 4°C. For the High-Salt Sarkosyl step, the sample was rotated for an additional 1 hour at room temperature prior to centrifugation. The sarkosyl insoluble pellet was sonicated (Qsonica) for 20 pulses at 100% amplitude with 0.5 s intervals in urea buffer before being centrifuged at 25,000 x g for 30 min at 4°C.

### Immunoprecipitation and mass spectrometry of sarkosyl brain fractions

The sarkosyl fractions from 3 control and 3 MSA pons were sonicated (Qsonica) for 20 pulses at 100% amplitude with 0.5 s intervals in DPBS, then immunoprecipitated using an anti-TPPP antibody (ThermoFisher PA5-19243) and Protein-G-conjugated Dynabeads (ThermoFisher) per manufacturer’s protocol. Beads were eluted by heating at 95°C for 5 min in 0.2% RapiGest (Waters) with 5 mM DTT and 50 mM ammonium bicarbonate.

For mass spectrometry, eluted in-solution samples were alkylated with N-Ethylmaleimide and digested with trypsin (Promega) using micro S-trap column (ProtiFi). Digests were desalted with an Oasis HLB µElution plate (Waters) then injected into the Ultimate 3000 HPLC system. An ES902 column (ThermoFisher) was used to separate the peptides at a flow rate of 300 nL/min. Mobile phase B (0.1% formic acid in ACN) amount was increased from 3% to 18% in 48 min, from 24% to 36% in 8 min, then followed by wash steps.

The LC-MS/MS data were acquired on a Fusion Lumos Orbitrap Mass Spectrometer (ThermoFisher) using the ETciD method. The precursor ion intensity threshold to trigger the MS/MS scan was set at 1 x 104. The quadrupole isolation window was 2 m/z. The MS2 scans were acquired in ion trap. MS1 scans were performed every 3 sec. As many MS2 scans were acquired within the MS1 scan cycle.

Proteome Discoverer software version 2.4 was used for data analysis. Raw data were searched against the Sprot Human database. Maximum allowed missed cleavage site was 3. Mass tolerances for MS1 and MS2 scans were set to 5 ppm and 0.6 Da, respectively. The search results were filtered by a false discovery rate of 1% at the protein level. Hits were filtered for those with p-values <0.05 (calculated from ANOVA), then sorted by normalized protein abundance ratios. For each sample, protein abundance values were calculated by summing the abundance of unique and razor peptides matched to that protein. Normalization was performed against the total peptide amount. Protein ratios were calculated by comparing the median protein abundances between control and MSA samples.

### Western blots

Samples were run on Mini-Protean TGX gels (BioRad), then transferred to PVDF membranes. Membranes were incubated with primary and HRP-conjugated secondary antibodies, then developed using SuperSignal West Pico PLUS substrate (ThermoFisher Scientific). For sequential extractions, each fraction was first quantified by BCA analysis to determine protein concentration, then 10 μg of denatured protein was loaded. Due to a large number of lanes, each gel was run with samples from 1 control and 1 MSA subject. The resulting blots were probed with antibodies for aSyn (Abcam ab209420, 1:1000), GAPDH (Proteintech 60004-1-Ig, 1:50,000), and TPPP (ThermoFisher PA5-19243, 1:1000), then imaged using an Azure Imaging Box. Western blot bands were quantified using the Analyze Gel function of Fiji.

### Immunopanning and culturing of primary oligodendrocytes

OPCs were purified from Sprague-Dawley rat pups (P6–P8) by immunopanning as previously described (*24*). Briefly, cortical tissue was dissociated by papain digestion then filtered through a Nitex mesh to obtain a mixed single-cell suspension. This suspension was incubated in 2 negative-selection plates coated with anti-Ran-2 and anti-GC antibodies, then in a positive-selection plate coated with anti-O4 antibody. Adherent cells were trypsinized and cultured in proliferation media containing PDGF and NT-3 or differentiation media containing T3.

For 3D cultures, 2-µm diameter electrospun microfibers (Amsbio, Mimetix aligned 12-well inserts) that mimic the shape of axons were coated with poly-D-lysine and immunopanned OPCs were seeded onto the microfibers in differentiation media.

### Primary hippocampal neuron culture

Hippocampal neurons from E18 rat embryos were isolated as described in the Lonza Amaxa Rat Neuron Nucleofector protocol and cultured for 15 days in Neurobasal media with 2 mM L-glutamine and 2% B-27 supplement.

### Oligodendrocyte live-cell imaging and FRAP analysis

Immunopanned OPCs were electroporated (Lonza Amaxa II, O-17 program) with a TPPP-EGFP construct, then plated onto No. 1.5 glass-bottom dishes coated with poly-D-lysine (MatTek). Cells were imaged on a Nikon Ti2 spinning-disk confocal microscope with Perfect Focus using a Hamamatsu ORCA FusionBT scMOS camera. Live-cell imaging was performed in an environmental chamber at 37°C with 5% CO_2_ (Tokai) while engaging the Perfect Focus unit of the microscope. To decrease phototoxicity, cells were imaged in differentiation media in which DMEM was replaced with FluoroBrite DMEM. Each dish was imaged for a total duration of 1 hour or less.

FRAP assays using the OMS Photostimulation System were performed using the JOBS module of the Nikon Elements software. To perform the bleach, small circular ROIs were created around a region containing TPPP-EGFP puncta in a cell process. A 405 laser line was used to target the ROI for the bleach stimulation event. The cell was imaged prior to the bleach event and then every 5 s immediately following the bleach event for over 1 minute. ROI fluorescence intensity data were collected and a uniform background subtraction was applied to all images. Analysis of the recovery curve and the determination of half-time was performed by fitting to a one phase exponential association curve using GraphPad Prism.

### Oligodendrocyte immunostaining

Oligodendrocytes were fixed in 4% paraformaldehyde, permeabilized using 0.1% Triton-X100, and blocked in 5% donkey serum with 1% BSA (bovine serum albumin). Cells were then stained using primary antibodies overnight at 4°C: tubulin (Invitrogen MA1-80017, 1:1000), aSyn (BD Transduction Laboratories 610786, 1:500), MBP (Abcam ab7349, 1:1000). Fluorescent dye conjugated donkey secondary antibodies (Jackson ImmunoResearch, 1:500) were applied for one hour at room temperature, followed by DAPI (Sigma). Micrographs were acquired on a Nikon Ti2 spinning-disk confocal using 60X or 100X objectives with a scMOS Hamamatsu ORCA-FusionBT camera.

### Tubulin and TPPP-EGFP colocalization

The line selection tool in Fiji was used to create 2-μm lines perpendicular to the oligodendrocyte process and the Plot Profile function was used to obtain the fluorescence intensity values of each channel. The data were normalized to the maximum and minimum intensity of each line-scan and plotted as separate graphs for each line.

### Constructs

For *in vitro* experiments, full length human TPPP (BC131506) and αSynuclein (MJFF Addgene 51486) were cloned into modified pHAT bacterial expression vectors as previously described (Fu et al. 2019). All vectors contain an N-terminal 6x-His-tag, C terminal Strep-Tag II tag, with or without a N or C terminal fluorophore (EGFP, mNeonGreen (Allele Biotech), or mRuby2).

### Recombinant protein expression

Recombinant protein was expressed using BL21(DE3) *E. coli* transformed with protein expression vectors. *E. coli* were grown at 37°C for 4–6 hours until they reached OD 0.4– 0.6, then they were cooled on ice until they reached 18°C. Expression was induced with 0.5 mM IPTG and cells were further incubated overnight at 18°C for ∼16 hours. Bacterial pellets were harvested with centrifugation.

For liquid condensate and nucleation assays, bacterial pellets were resuspended in 25 mL Buffer A (50 mM Na2HPO4, 300 mM NaCl, pH 7.2) and were stored at −80°C until used for purification. Cells were first lysed using an EmulsiFlex-C5 (Avestin) and supernatant was clarified with centrifugation. Clarified supernatant was incubated with His60 NTA resin (TAKARA Bio) for 1 hour then loaded onto gravity flow columns and eluted with Buffer A containing 200 mM Imidazole. Proteins were further purified by loading His60 eluate onto Strep-TactinXT affinity chromatography gravity columns (IBA Lifesciences, Germany). Final pure protein was eluted with Buffer BXT +10% glycerol (IBA Lifesciences) or BRB80 (80 mM PIPES-KOH, 1 mM EGTA, 1mM MgCl_2_, pH 6.8) +50 mM D-biotin +10% glycerol.

For PFFs, bacterial pellets were resuspended in RLS Buffer (1 M KCl, 50 mM PIPES pH 6.8, and 2 mM DTT) supplied with EDTA-free protease inhibitor cocktail (Roche). The lysis was performed using homogenizer EmulsiFlex-C3 (Avestin). The lysate was separated into soluble and insoluble fractions by ultracentrifugation at 100,000 x g for 1 h using Optima L-XP (Beckman Coulter). The soluble fraction was filtered using a bottle-top 0.22-µm filter (Corning). The protein was purified from the soluble fraction by sequential affinity purification in gravity flow columns (Bio-Rad) using His60 Ni Superflow Resin (Qiagen) then using Strep-Tactin®XT 4Flow® high capacity resin (IBA) with biotin elution in RLS Buffer.

Tubulin was purified from bovine brains as previously described (*46*) with the modification of using Fractogel EMD SO_3_^−^(M) resin (Millipore-Sigma) instead of phosphocellulose. Tubulin was labeled using Atto633-NHS ester (ATTO-TEC) or CF640R (Biotium) as previously described (*47*). An additional cycle of polymerization/depolymerization was performed before use.

Proteins were used fresh for experiments or flash frozen and stored in liquid nitrogen. Protein concentrations were determined at 280 nm using a DS-11 FX spectrophotometer (DeNovix), then calculated using the Beer–Lambert Law. Purity of all protein reagents was determined using SDS-PAGE and/or MS.

### Glass and imaging chamber preparation

Cover glasses were prepared by washing in acetone for 10 min, then sonicating in 50% methanol for 20 min, then sonicating in 0.5 M KOH for 20 min, then washing 3 times with water. Coverslips were then dried with nitrogen gas before being exposed to air plasma (Plasma Etch) for 3 min, then silanized by soaking in 0.2% Dichlorodimethylsilane (DDS) in n-Heptane for 2 hours. Coverslips were finished by sonication in n-Heptane for 20 min, then sonication in 100% ethanol for 20 min, then rinsed with water and dried with nitrogen gas. Flow channels were constructed using two pieces of silanized cover glasses (22 X 22 mm and 18 X 18 mm) held together with double-sided tape and mounted into custom-machined coverslip holders. Glass surfaces were blocked with 5% F-127 in BRB80 before use.

### Phase separation and FRAP assays

For characterization of TPPP condensates, droplets were formed with EGFP-TPPP, TPPP-mNeonGreen (mNG), or mRuby2-TPPP. For heterogeneous condensate experiments, we added 6% labeled tubulin (Atto-633 or CF640R) and/or αSynuclein-mNG. All experiments were conducted in imaging buffer (BRB80 supplemented with 0.1 mg/mL BSA, 10mM DTT, 250 nM glucose oxidase, 64 nM catalase, 40 mM D-glucose) with varying concentrations of KCl (0–2000 mM). Some experiments were conducted with 10% (w/v) PEG 8000 (Sigma) as indicated in the figure legends.

For droplet fusion experiments, images were taken at 100 ms intervals. The ability of GPF-TPPP to form droplets was tested under increasing KCl concentration (0– 2000 nM). For TPPP surface wetting images, F-127 was not used to block the glass surface.

For FRAP, TPPP and tubulin condensates were bleached with a 488 nM or 637 nM laser (100% laser power) for ∼10 s using iLAS2 laser illumination (BioVision). Before and after bleaching, images were taken at ∼0.1–0.4 s intervals with ∼50–100 ms exposure and 6% laser power. Analysis of the recovery curves and the determination of half-time recovery was performed by fitting to a single exponential y(t)=A(1-e^-t*x^)+c in Python 3 (python.org) using a JupyterLab Notebook.

### Microtubule nucleation assays

Microtubule nucleation from TPPP condensates was visualized using TIRF microscopy. Flow channels were first incubated with β3-tubulin antibody (BioLegend:Poly18020) 1:100 in BRB80 then blocked with 5% F-127 in BRB80. Nucleation was achieved by flowing in 200–1000 nM EGFP-TPPP with 8 µM tubulin (6% CF640R) in imaging buffer with 1 mM GTP. Microtubule nucleation was also imaged with unlabeled TPPP and in the absence of PEG.

For analysis of microtubule nucleation, condensates were thresholded in the TPPP channel to include the region at the center of microtubule nucleation, but not include any protruding microtubules. This thresholded region was then used to calculate area and integrated density (area*mean signal) of both TPPP and tubulin.

### *In vitro* microscopy

For FRAP experiments, images were recorded with a Zeiss Axiovert Z1 microscope chassis and a 100 ×1.45 NA Plan-apochromat objective lens. TIRF was achieved by coupling 488/561/637 nm lasers to an iLas2 targeted laser illumination system (BioVision, Inc) equipped with 360° TIRF. The objective was heated to 35°C with a CU-501 Chamlide lens warmer (Live Cell Instrument). For microtubule nucleation experiments data was acquired with a customized Zeiss Axio Observer seven equipped with a Laser TIRF III and 405/488/561/638 nm lasers, Alpha Plan-Apo 100 ×/1.46Oil DIC M27, and Objective Heater 25.5/33 S1. Images on both systems were recorded on a Prime 95B CMOS camera (Photometrics) with a pixel size of 107 nm.

### Image analysis and software (*in vitro*)

Partition coefficients were determined by measuring the mean signal inside droplets divided by the background outside the droplet. All *in vitro* images were processed and analyzed using Fiji (*45*). Images were linearly adjusted in brightness and contrast using Photoshop (Adobe).

### TPPP PFF formation and analysis

Purified TPPP was dialyzed overnight in 0.5–3 mL dialysis cassettes (Thermo Scientific) against PBS. Soluble protein was pre-cleared by ultracentrifugation at 100,000 x g for 1 hour, then diluted with PBS to a final concentration of 7 µM in low-binding tubes. For PFF formation, TPPP was incubated in a benchtop shaking heat block at 37°C with agitation at 1,000 RPM for 48 hours.

PFFs were fragmented using a sonicator (Qsonica) set to amplitude 100% for 60 pulses with duration of 0.5 s, interspersed with 10-s breaks between every 10 pulses. Sonicated PFFs were snap-frozen in liquid nitrogen and stored in liquid nitrogen. The absorbance of TPPP was measured at wavelength 280 nm using a spectrophotometer.

### PFF TEM and immunogold labeling

Negative staining was performed for analysis of PFF morphology. Briefly, 3 µL of PFFs was applied onto carbon-coated 300-mesh copper grids (EMS). The grids were stained with 2% uranyl acetate (EMS).

Immunogold labeling was performed by applying TPPP PFFs to carbon-coated 400-mesh nickel grids (SPI). Grids were incubated with an anti-TPPP primary antibody (ProteinTech, rabbit, diluted 1:100) for 30 min at room temperature. The grid was subsequently washed 3 times with PBS, then incubated in a secondary goat anti-rabbit antibody conjugated to 10-nm colloidal gold (Ted Pella, diluted 1:100) for 30 min at room temperature. After washing with PBS, the sample was fixed with 8% glutaraldehyde (EMS) for 5 min and negative staining was performed with 1% uranyl acetate (EMS) for 1 min.

All PFF imaging was performed using JEOL JEM-200CX at high voltages of 80– 120 kV. Quantification of number and size of TPPP PFFs was performed using the pixel and object classification functions of the machine-learning software ilastik (*44*). Specificity of immunogold labeling was determined by calculating the number of gold beads per area using the freehand selection tool and measure function in Fiji (41).

### Oligodendrocyte PFF toxicity assays

Immunopanned rat oligodendrocytes were incubated with sonicated TPPP PFFs. For live-cell imaging assays, annexin V green dye (Sartorius cat. #4642, 1:200) and a final concentration of 0.07uM TPPP PFFs were added to imaging media made with Fluorobrite DMEM (Fisher cat. #A18967-01). DIV 1 oligodendrocytes were imaged using a 10X objective across ∼80–100 stitched fields of view every 1 hour for 5 days. Half the media was changed every 2 days to refresh the TPPP PFF’s and annexin dye. Quantification of annexin positive structures was done using Nikon Elements General Analysis. Total area analyzed was 29.03 mm^2^ (control) and 30.42 mm^2^ (TPPP treated), not including the densely plated central region of the coverslip which did not allow for distinguishing individual structures and also exhibited high levels of normal cell death that occurs during OPC differentiation in a dense cellular environment *in vitro*.

For TUNEL analysis, DIV4 oligodendrocytes were incubated with either vehicle (PBS) or 0.7 µM, 0.07 µM, or 0.007 µM TPPP PFFs. After 24 hours, cells were fixed and stained using the Click-iT Plus TUNEL assay kit (Invitrogen C10619) and DAPI and immunostained for tubulin (Sigma T9026, 1:1000). DAPI and TUNEL quantification was done using Nikon Elements General Analysis and confirmed manually (investigator was blinded to experimental conditions).

### Statistics

Means, standard deviations, standard errors, and statistical analyses were calculated and graphed using OriginPro2020 (OriginLab) or Prism 9 (GraphPad). Data were first tested for normality or parametricity using Shapiro-Wilk and Kolmogorov-Smirnov tests. Normal data was analyzed using t-test or one-way ANOVA. Non-normal data was analyzed using the Kruskal-Wallis test.

## ACKNOWLEDGEMENTS

We thank the NINDS (National Institute of Neurological Disorders and Stroke) Electron Microscopy Core (Susan Cheng, Sandra Moreira) for training and assistance with TEM. We thank Boma Fubara (NINDS mass spectroscopy core) for technical assistance. We thank the following NINDS labs for use of their equipment/reagents, technical training, purchasing assistance, and mentorship: Antonina Roll-Mecak Lab (Kishore Mahalingan), Kenton Swartz Lab (Helena Tsg-Hui Chang), Joseph Mindell Lab (Vedrana Mikusevic), and Miguel Holmgren (Deepa Srikumar). We thank Derek Narendra for recommending a TH antibody. We thank the Harvard Brain Tissue Resource Center (Sabina Berretta) via the NIH NeuroBioBank for providing human brain samples. Finally, we are grateful to the patients who donated their brains for their final and lasting contributions. M.-m.F. is supported by NINDS intramural funding (NS009432). H.S.R. is partially supported by the NINDS Intramural Diversity Training Supplement program. S.B. is supported by NSERC RGPIN-2017-04649, CIHR PJT-156193, and a grant from the Defeat MSA Alliance. T.S.M is supported by Fonds de Recherche du Québec - Santé (Doctoral Training).

## AUTHOR CONTRIBUTIONS

Conceptualization, M.-m.F., S.B., S.K.; Methodology, M.-m.F., S.B., S.K., H.S.R., T.S.M., A.K., J.C.N., Y.L.; Formal analysis, M.-m.F., S.B., S.K., H.S.R., T.S.M., A.K., J.C.N., Y.L.; Investigation, S.K., H.S.R., T.S.M., A.K., J.C.N., M.-m.F., S.B.; Writing – Original Draft, S.K., H.S.R., T.S.M., A.K., M.-m. F, S.B.; Writing – Review & Editing, S.K., H.S.R., M.-m. F, S.B.; Visualization, S.K., H.S.R., T.S.M., A.K., J.C.N., M.-m.F.,S.B.; Supervision, M.-m.F., S.B.; Funding Acquisition, M.-m.F., S.B

## MATERIALS & CORRESPONDENCE

Correspondence and material requests should be addressed to Meng-meng Fu.

## REFERENCES

1. M. Weigel, L. Wang, M. Fu, Microtubule organization and dynamics in oligodendrocytes, astrocytes, and microglia. Dev. Neurobiol. 81(3), 310–320 (2021).

2. A. L. Herbert, M. M. Fu, C. M. Drerup, R. S. Gray, B. L. Harty, S. D. Ackerman, T. O’Reilly-Pol, S. L. Johnson, A. V. Nechiporuk, B. A. Barres, K. R. Monk, Dynein/dynactin is necessary for anterograde transport of Mbp mRNA in oligodendrocytes and for myelination in vivo. Proc Natl Acad Sci U A. 114, E9153–E9162 (2017).

3. N. Snaidero, W. Mobius, T. Czopka, L. H. Hekking, C. Mathisen, D. Verkleij, S. Goebbels, J. Edgar, D. Merkler, D. A. Lyons, K. A. Nave, M. Simons, Myelin membrane wrapping of CNS axons by PI(3,4,5)P3-dependent polarized growth at the inner tongue. Cell. 156, 277–90 (2014).

4. J. M. Edgar, E. McGowan, K. J. Chapple, W. Möbius, L. Lemgruber, R. H. Insall, K. Nave, A. Boullerne, RíoLHortega’s drawings revisited with fluorescent protein defines a cytoplasmLfilled channel system of CNS myelin. J. Anat. 239, 1241–1255 (2021).

5. M. M. Fu, T. S. McAlear, H. Nguyen, J. A. Oses-Prieto, A. Valenzuela, R. D. Shi, J. J. Perrino, T. T. Huang, A. L. Burlingame, S. Bechstedt, B. A. Barres, The Golgi Outpost Protein TPPP Nucleates Microtubules and Is Critical for Myelination. Cell. 179, 132–146 e14 (2019).

6. A. Valenzuela, L. Meservey, H. Nguyen, M. Fu, Golgi Outposts Nucleate Microtubules in Cells with Specialized Shapes. Trends Cell Biol., S0962892420301458 (2020).

7. H. Nguyen, L. M. Meservey, N. Ishiko-Silveria, M. Zhou, T.-T. Huang, M. Fu, Fear Deficits in Hypomyelinated *Tppp* Knock-Out Mice. eNeuro. 7(5) (2020).

8. G. G. Kovacs, L. Laszlo, J. Kovacs, P. H. Jensen, E. Lindersson, G. Botond, T. Molnar, A. Perczel, F. Hudecz, G. Mezo, A. Erdei, L. Tirian, A. Lehotzky, E. Gelpi, H. Budka, J. Ovadi, Natively unfolded tubulin polymerization promoting protein TPPP/p25 is a common marker of alpha-synucleinopathies. Neurobiol Dis. 17, 155–62 (2004).

9. E. Lindersson, D. Lundvig, C. Petersen, P. Madsen, J. R. Nyengaard, P. Hojrup, T. Moos, D. Otzen, W. P. Gai, P. C. Blumbergs, P. H. Jensen, p25alpha Stimulates alpha-synuclein aggregation and is co-localized with aggregated alpha-synuclein in alpha-synucleinopathies. J Biol Chem. 280, 5703–15 (2005).

10. Y. Zhang, K. Chen, S. A. Sloan, M. L. Bennett, A. R. Scholze, S. O’Keeffe, H. P. Phatnani, P. Guarnieri, C. Caneda, N. Ruderisch, S. Deng, S. A. Liddelow, C. Zhang, R. Daneman, T. Maniatis, B. A. Barres, J. Q. Wu, An RNA-sequencing transcriptome and splicing database of glia, neurons, and vascular cells of the cerebral cortex. J Neurosci. 34, 11929– 47 (2014).

11. Q. R. Lu, D. Yuk, J. A. Alberta, Z. Zhu, I. Pawlitzky, J. Chan, A. P. McMahon, C. D. Stiles, D. H. Rowitch, Sonic hedgehog--regulated oligodendrocyte lineage genes encoding bHLH proteins in the mammalian central nervous system. Neuron. 25, 317–329 (2000).

12. M. Neumann, D. M. Sampathu, L. K. Kwong, A. C. Truax, M. C. Micsenyi, T. T. Chou, J. Bruce, T. Schuck, M. Grossman, C. M. Clark, L. F. McCluskey, B. L. Miller, E. Masliah, I. R. Mackenzie, H. Feldman, W. Feiden, H. A. Kretzschmar, J. Q. Trojanowski, V. M.-Y. Lee, Ubiquitinated TDP-43 in Frontotemporal Lobar Degeneration and Amyotrophic Lateral Sclerosis. Science. 314, 130–133 (2006).

13. A. Molliex, J. Temirov, J. Lee, M. Coughlin, A. P. Kanagaraj, H. J. Kim, T. Mittag, J. P. Taylor, Phase separation by low complexity domains promotes stress granule assembly and drives pathological fibrillization. Cell. 163, 123–33 (2015).

14. H. B. Schmidt, R. Rohatgi, In Vivo Formation of Vacuolated Multi-phase Compartments Lacking Membranes. Cell Rep. 16, 1228–1236 (2016).

15. A. Patel, H. O. Lee, L. Jawerth, S. Maharana, M. Jahnel, M. Y. Hein, S. Stoynov, J. Mahamid, S. Saha, T. M. Franzmann, A. Pozniakovski, I. Poser, N. Maghelli, L. A. Royer, M. Weigert, E. W. Myers, S. Grill, D. Drechsel, A. A. Hyman, S. Alberti, A Liquid-to-Solid Phase Transition of the ALS Protein FUS Accelerated by Disease Mutation. Cell. 162, 1066–77 (2015).

16. T. Murakami, S. Qamar, J. Q. Lin, G. S. Schierle, E. Rees, A. Miyashita, A. R. Costa, R. B. Dodd, F. T. Chan, C. H. Michel, D. Kronenberg-Versteeg, Y. Li, S. P. Yang, Y. Wakutani, W. Meadows, R. R. Ferry, L. Dong, G. G. Tartaglia, G. Favrin, W. L. Lin, D. W. Dickson, M. Zhen, D. Ron, G. Schmitt-Ulms, P. E. Fraser, N. A. Shneider, C. Holt, M. Vendruscolo, C. F. Kaminski, P. St George-Hyslop, ALS/FTD Mutation-Induced Phase Transition of FUS Liquid Droplets and Reversible Hydrogels into Irreversible Hydrogels Impairs RNP Granule Function. Neuron. 88, 678–90 (2015).

17. D. T. Jones, D. Cozzetto, DISOPRED3: precise disordered region predictions with annotated protein-binding activity. Bioinforma. Oxf. Engl. 31, 857–863 (2015).

18. M. J. Mizianty, Z. Peng, L. Kurgan, MFDp2: Accurate predictor of disorder in proteins by fusion of disorder probabilities, content and profiles. Intrinsically Disord. Proteins. 1, e24428 (2013).

19. T. Ishida, K. Kinoshita, PrDOS: prediction of disordered protein regions from amino acid sequence. Nucleic Acids Res. 35, W460–464 (2007).

20. M. Hardenberg, A. Horvath, V. Ambrus, M. Fuxreiter, M. Vendruscolo, Widespread occurrence of the droplet state of proteins in the human proteome. Proc. Natl. Acad. Sci. U. S. A. 117, 33254–33262 (2020).

21. J. B. Woodruff, B. Ferreira Gomes, P. O. Widlund, J. Mahamid, A. Honigmann, A. A. Hyman, The Centrosome Is a Selective Condensate that Nucleates Microtubules by Concentrating Tubulin. Cell. 169, 1066–1077 e10 (2017).

22. S. Petry, A. C. Groen, K. Ishihara, T. J. Mitchison, R. D. Vale, Branching microtubule nucleation in Xenopus egg extracts mediated by augmin and TPX2. Cell. 152, 768–77 (2013).

23. T. Consolati, J. Locke, J. Roostalu, Z. A. Chen, J. Gannon, J. Asthana, W. M. Lim, F. Martino, M. A. Cvetkovic, J. Rappsilber, A. Costa, T. Surrey, Microtubule Nucleation Properties of Single Human γTuRCs Explained by Their Cryo-EM Structure. Dev. Cell. 53, 603–617.e8 (2020).

24. J. C. Dugas, B. Emery, Purification of oligodendrocyte precursor cells from rat cortices by immunopanning. Cold Spring Harb Protoc. 2013, 745–58 (2013).

25. V.-I. Gargareta, J. Reuschenbach, S. B. Siems, T. Sun, L. Piepkorn, C. Mangana, E. Späte, S. Goebbels, I. Huitinga, W. Möbius, K.-A. Nave, O. Jahn, H. B. Werner, Conservation and divergence of myelin proteome and oligodendrocyte transcriptome profiles between humans and mice. eLife. 11, e77019 (2022).

26. J. Burré, M. Sharma, T. Tsetsenis, V. Buchman, M. R. Etherton, T. C. Südhof, Alpha-synuclein promotes SNARE-complex assembly in vivo and in vitro. Science. 329, 1663– 1667 (2010).

27. T. Logan, J. Bendor, C. Toupin, K. Thorn, R. H. Edwards, α-Synuclein promotes dilation of the exocytotic fusion pore. Nat. Neurosci. 20, 681–689 (2017).

28. M. Djelloul, S. Holmqvist, A. Boza-Serrano, C. Azevedo, M. S. Yeung, S. Goldwurm, J. Frisén, T. Deierborg, L. Roybon, Alpha-Synuclein Expression in the Oligodendrocyte Lineage: an In Vitro and In Vivo Study Using Rodent and Human Models. Stem Cell Rep. 5, 174–184 (2015).

29. Y. Zhang, S. A. Sloan, L. E. Clarke, C. Caneda, C. A. Plaza, P. D. Blumenthal, H. Vogel, G. K. Steinberg, M. S. Edwards, G. Li, J. A. Duncan 3rd, S. H. Cheshier, L. M. Shuer, E. F. Chang, G. A. Grant, M. G. Gephart, B. A. Barres, Purification and Characterization of Progenitor and Mature Human Astrocytes Reveals Transcriptional and Functional Differences with Mouse. Neuron. 89, 37–53 (2016).

30. S. Ray, N. Singh, R. Kumar, K. Patel, S. Pandey, D. Datta, J. Mahato, R. Panigrahi, A. Navalkar, S. Mehra, L. Gadhe, D. Chatterjee, A. S. Sawner, S. Maiti, S. Bhatia, J. A. Gerez, A. Chowdhury, A. Kumar, R. Padinhateeri, R. Riek, G. Krishnamoorthy, S. K. Maji, α-Synuclein aggregation nucleates through liquid-liquid phase separation. Nat. Chem. 12, 705–716 (2020).

31. C. Peng, R. J. Gathagan, D. J. Covell, C. Medellin, A. Stieber, J. L. Robinson, B. Zhang, R. M. Pitkin, M. F. Olufemi, K. C. Luk, J. Q. Trojanowski, V. M.-Y. Lee, Cellular milieu imparts distinct pathological α-synuclein strains in α-synucleinopathies. Nature. 557, 558–563 (2018).

32. I. V. J. Murray, B. I. Giasson, S. M. Quinn, V. Koppaka, P. H. Axelsen, H. Ischiropoulos, J. Q. Trojanowski, V. M.-Y. Lee, Role of alpha-synuclein carboxy-terminus on fibril formation in vitro. Biochemistry. 42, 8530–8540 (2003).

33. K. C. Luk, C. Song, P. O’Brien, A. Stieber, J. R. Branch, K. R. Brunden, J. Q. Trojanowski, V. M.-Y. Lee, Exogenous alpha-synuclein fibrils seed the formation of Lewy body-like intracellular inclusions in cultured cells. Proc. Natl. Acad. Sci. U. S. A. 106, 20051–20056 (2009).

34. L. A. Volpicelli-Daley, K. C. Luk, T. P. Patel, S. A. Tanik, D. M. Riddle, A. Stieber, D. F. Meaney, J. Q. Trojanowski, V. M.-Y. Lee, Exogenous α-synuclein fibrils induce Lewy body pathology leading to synaptic dysfunction and neuron death. Neuron. 72, 57–71 (2011).

35. L. O. Sun, S. B. Mulinyawe, H. Y. Collins, A. Ibrahim, Q. Li, D. J. Simon, M. Tessier-Lavigne, B. A. Barres, Spatiotemporal Control of CNS Myelination by Oligodendrocyte Programmed Cell Death through the TFEB-PUMA Axis. Cell. 175, 1811–1826.e21 (2018).

36. S. E. Logue, M. Elgendy, S. J. Martin, Expression, purification and use of recombinant annexin V for the detection of apoptotic cells. Nat. Protoc. 4, 1383–1395 (2009).

37. M. R. King, S. Petry, Phase separation of TPX2 enhances and spatially coordinates microtubule nucleation. Nat. Commun. 11, 270 (2020).

38. W. T. Snead, A. S. Gladfelter, The Control Centers of Biomolecular Phase Separation: How Membrane Surfaces, PTMs, and Active Processes Regulate Condensation. Mol. Cell. 76, 295–305 (2019).

39. M. Takahashi, K. Tomizawa, S. C. Fujita, K. Sato, T. Uchida, K. Imahori, A Brain-Specific Protein p25 Is Localized and Associated with Oligodendrocytes, Neuropil, and Fiber-Like Structures of the CA3 Hippocampal Region in the Rat Brain. J. Neurochem. 60, 228–235 (1993).

40. A. G. de la Fuente, R. M. L. Queiroz, T. Ghosh, C. E. McMurran, J. F. Cubillos, D. E. Bergles, D. C. Fitzgerald, C. A. Jones, K. S. Lilley, C. P. Glover, R. J. M. Franklin, Changes in the Oligodendrocyte Progenitor Cell Proteome with Ageing. Mol. Cell. Proteomics. 19, 1281–1302 (2020).

41. H. Mathys, J. Davila-Velderrain, Z. Peng, F. Gao, S. Mohammadi, J. Z. Young, M. Menon, L. He, F. Abdurrob, X. Jiang, A. J. Martorell, R. M. Ransohoff, B. P. Hafler, D. A. Bennett, M. Kellis, L.-H. Tsai, Single-cell transcriptomic analysis of Alzheimer’s disease. Nature. 570, 332–337 (2019).

42. S. Frykman, Y. Teranishi, J. Y. Hur, A. Sandebring, N. G. Yamamoto, M. Ancarcrona, T. Nishimura, B. Winblad, N. Bogdanovic, S. Schedin-Weiss, T. Kihara, L. O. Tjernberg, Identification of two novel synaptic gamma-secretase associated proteins that affect amyloid beta-peptide levels without altering Notch processing. Neurochem Int. 61, 108–18 (2012).

43. C. Simons, N. I. Wolf, N. McNeil, L. Caldovic, J. M. Devaney, A. Takanohashi, J. Crawford, K. Ru, S. M. Grimmond, D. Miller, D. Tonduti, J. L. Schmidt, R. S. Chudnow, R. van Coster, L. Lagae, J. Kisler, J. Sperner, M. S. van der Knaap, R. Schiffmann, R. J. Taft, A. Vanderver, A de novo mutation in the beta-tubulin gene TUBB4A results in the leukoencephalopathy hypomyelination with atrophy of the basal ganglia and cerebellum. Am J Hum Genet. 92, 767–73 (2013).

44. S. Berg, D. Kutra, T. Kroeger, C. N. Straehle, B. X. Kausler, C. Haubold, M. Schiegg, J. Ales, T. Beier, M. Rudy, K. Eren, J. I. Cervantes, B. Xu, F. Beuttenmueller, A. Wolny, C. Zhang, U. Koethe, F. A. Hamprecht, A. Kreshuk, ilastik: interactive machine learning for (bio)image analysis. Nat. Methods. 16, 1226–1232 (2019).

45. J. Schindelin, I. Arganda-Carreras, E. Frise, V. Kaynig, M. Longair, T. Pietzsch, S. Preibisch, C. Rueden, S. Saalfeld, B. Schmid, J. Y. Tinevez, D. J. White, V. Hartenstein, K. Eliceiri, P. Tomancak, A. Cardona, Fiji: an open-source platform for biological-image analysis. Nat Methods. 9, 676–82 (2012).

46. A. J. Ashford, A. A. Hyman, “Chapter 22 - Preparation of Tubulin from Porcine Brain” in Cell Biology, J. E. Celis, Ed. (Academic Press, Burlington, 2006), pp. 155–160.

47. A. Hyman, D. Drechsel, D. Kellogg, S. Salser, K. Sawin, P. Steffen, L. Wordeman, T. Mitchison, Preparation of modified tubulins. Methods Enzym. 196, 478–85 (1991).

