## Supplemental Data for "TPPP Forms Liquid Condensates and Aggregates in Multiple System Atrophy"

This document includes:

Figs. S1 to S6
Table S1
Titles for Movies S1 to S4

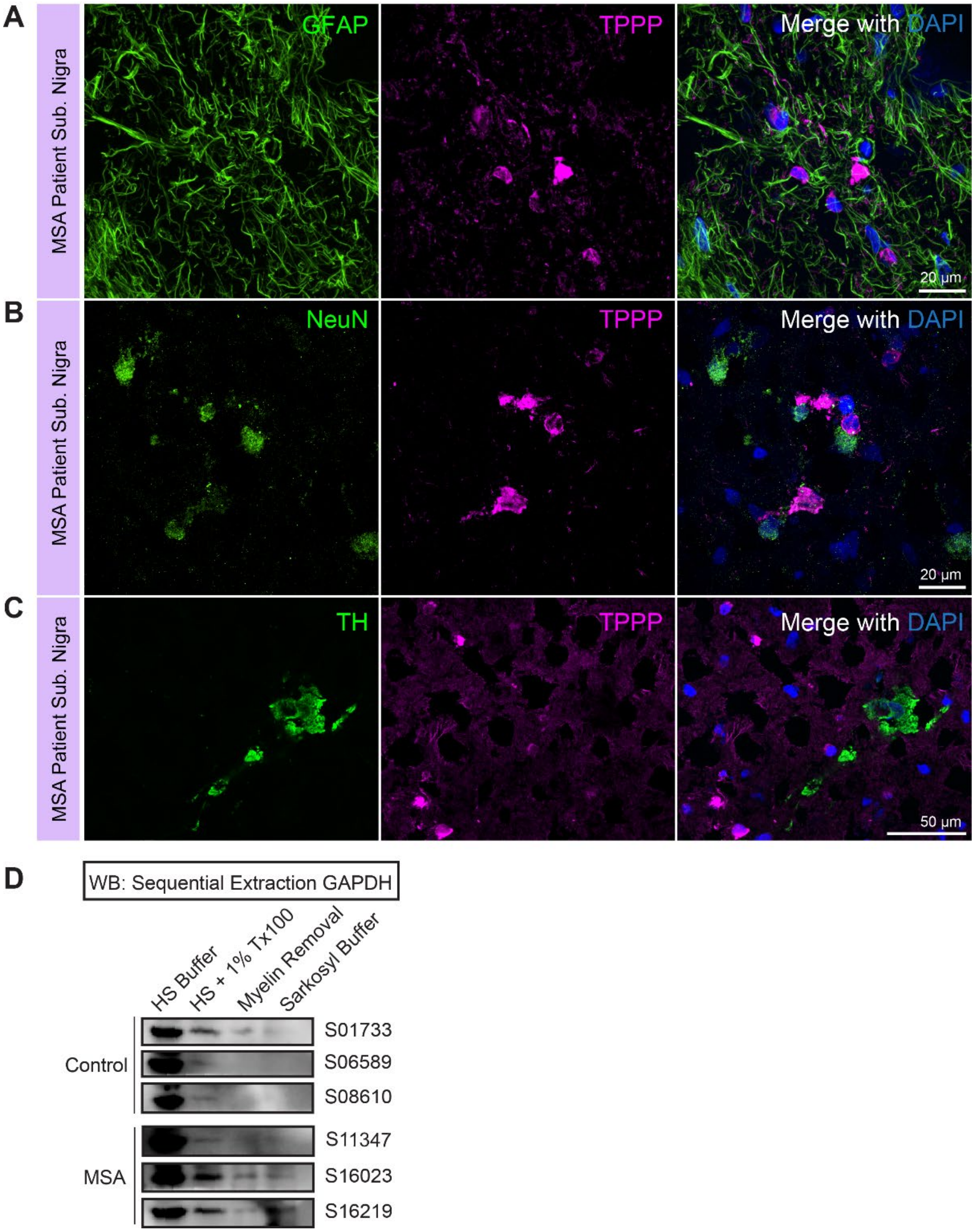


**Fig. S1. TPPP perinuclear staining does not colocalize with astrocyte or neuronal markers.**
(A) Micrographs of TPPP and the astrocyte marker GFAP in SN from MSA brain (Patient ID: S11347).
(B) Micrographs of TPPP and the neuronal marker NeuN in SN from MSA brain (Patient ID: S16219).
(C) Micrographs of TPPP and the dopaminergic neuron marker tyrosine hydroxylase (TH) in substantia nigra (SN) from MSA brain (Patient ID: S16219).
(D) Western blot of sequential extraction in increasingly stringent buffers in pons from control and MSA brains. Stained with GAPDH. n = 3 patients each control and MSA.

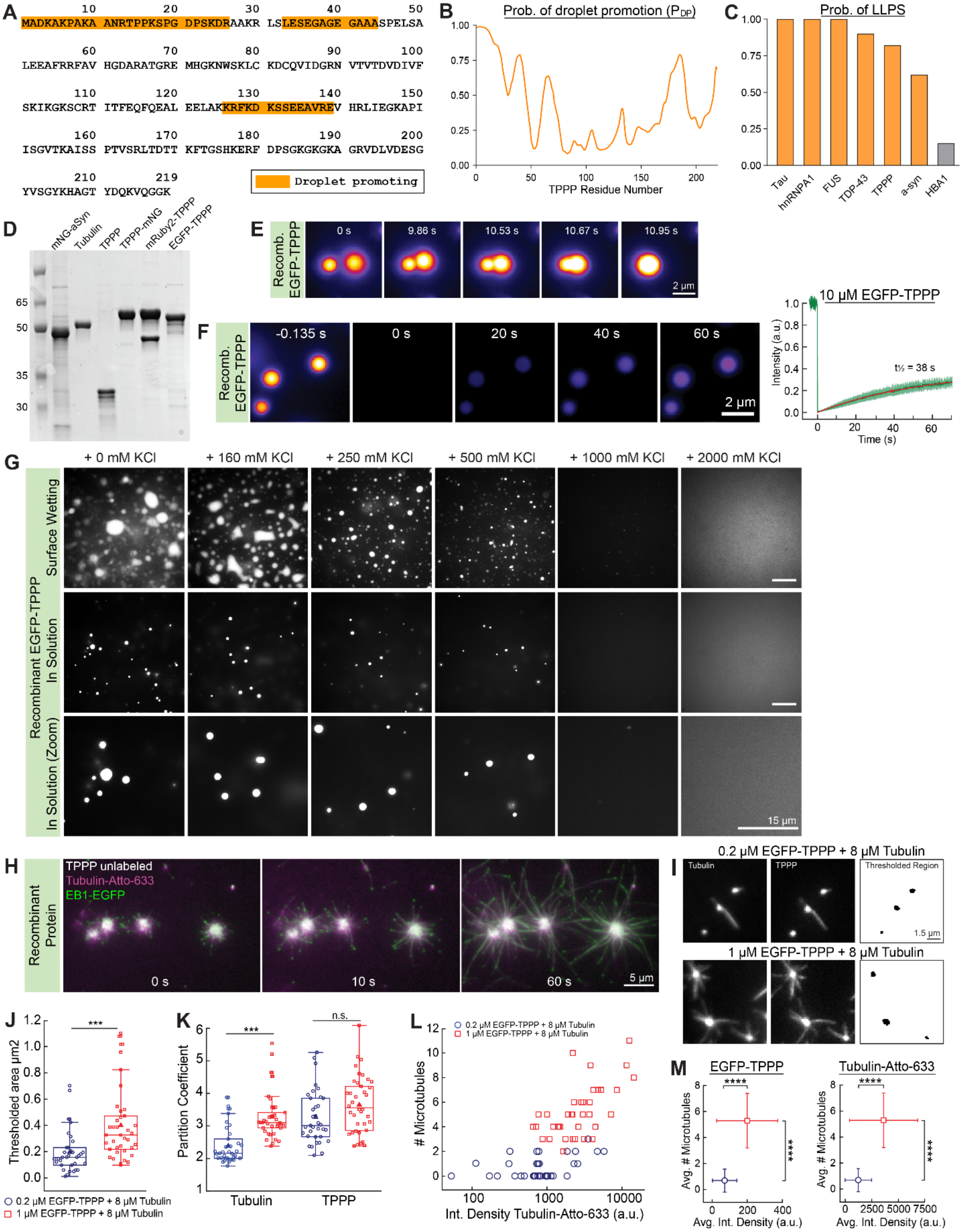


**Fig. S2. TPPP forms liquid condensates and nucleates microtubules.**
(A) Bioinformatics prediction of droplet-promoting regions for human TPPP using FuzDrop.

(B) FuzDrop probability of droplet promotion ($P_{DP}$) for human TPPP.
(C) FuzDrop probability of spontaneous LLPS ($P_{LLPS}$) for TPPP and other validated LLPS proteins and HBA1.
(D) SDS-PAGE gels of purified recombinant proteins. 1 µg each of mNG-aSyn, tubulin, TPPP, TPPP-mNG, mRuby2-TPPP, and EGFP-TPPP were loaded in each lane.
(E) Fast fusion of EGFP-TPPP condensates (10 µM) under fluid flow in BRB80 with 100 mM KCl and 10% PEG 8000. Images are pseudo-colored (yellow = high intensity; purple = low intensity).
(F) FRAP of 10 µM EGFP-TPPP and recovery curve with half-time of 38 s (BRB80 + 100 nM KCl + 10% PEG 8000). n = 5 condensates.
(G) EGFP-TPPP condensates (5 µM) in BRB80 with 10% PEG 8000 are inhibited with increasing concentrations of salt (0–2000 mM KCl). Condensates were imaged at the glass surface (top row), and in solution (middle row; bottom row enlarged images).
(H) Microtubule nucleation from TPPP condensates. Condensates were formed from 2 µM unlabeled TPPP, 15 µM Tubulin-Atto-633 and 250 nM EB1-EGFP in imaging buffer containing 1 mM GTP and 10% PEG 8000.
(I)TIRF images of TPPP condensates nucleating microtubules (0.2 µM or 1 µM EGFP-TPPP with 8 µM tubulin in BRB80 without PEG). Thresholded region (black) shows region used to calculate TPPP and tubulin puncta area and integrated density.
(J) Plot of thresholded area size of TPPP condensates. In Figs. S2J–S2M, blue circles represent 0.2 µM EGFP-TPPP + 8 µM tubulin (n = 35 condensates), red squares represent 1 µM EGFP-TPPP + 8 µM tubulin (n = 38 condensates). One-way ANOVA with post-hoc Tukey test.
(K) Partition coefficients of TPPP and tubulin condensates from microtubule nucleation experiments. One-way ANOVA with post-hoc Tukey test.
(L) Scatter plot of tubulin-CF640R condensate integrated density *vs*. number of microtubules nucleated.
(M) Plot of average integrated densities of EGFP-TPPP and tubulin-CF640R condensates *vs*. number of microtubules nucleated.

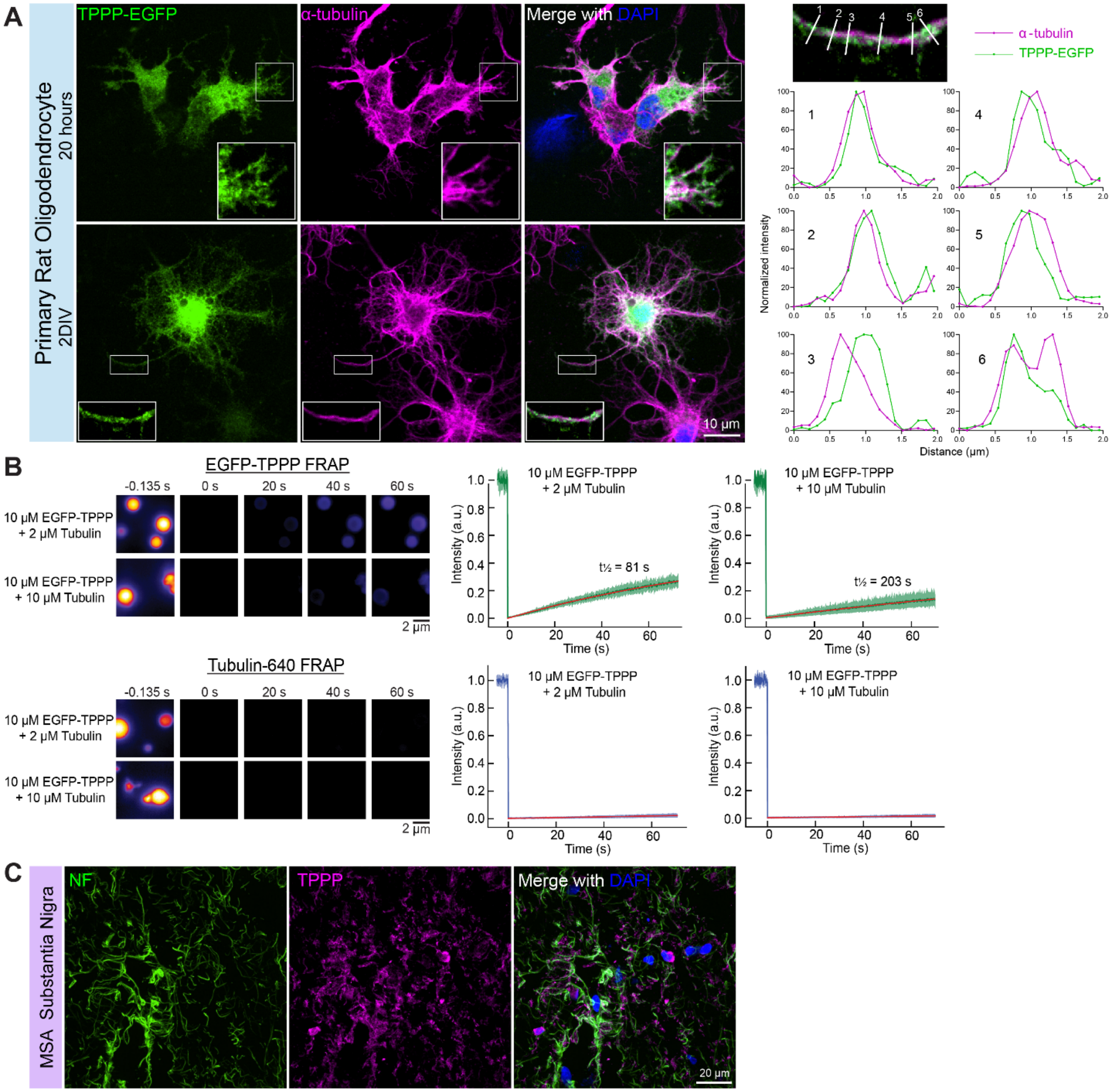


**Fig. S3. Filamentous/fibrillar TPPP found in primary oligodendrocytes and in MSA.**

(A) Oligodendrocytes expressing filamentous TPPP-EGFP for 20 hours and 2 days *in vitro* (DIV) stained against tubulin. Linescans of 2 DIV inset indicate that TPPP-EGFP forms clumps along microtubules.

(B) Representative immunofluorescent images and recovery plots of EGFP-TPPP or tubulin-CF640R before and after FRAP. Images are pseudo-colored (yellow = high intensity; purple = low intensity). Time of bleaching is defined as 0 s. Data are represented as mean ± SD (green) and fit to a single exponential $y(t)=A(1-e^{-t*t})+c$ (red). For EGFP-TPPP FRAP, n= 8, 5 condensates. For tubulin-CF640R, n = 6, 7 condensates.

(C) Micrographs of TPPP and the axonal marker neurofilament (NF) staining in SN from MSA brain (Patient ID: S11347).

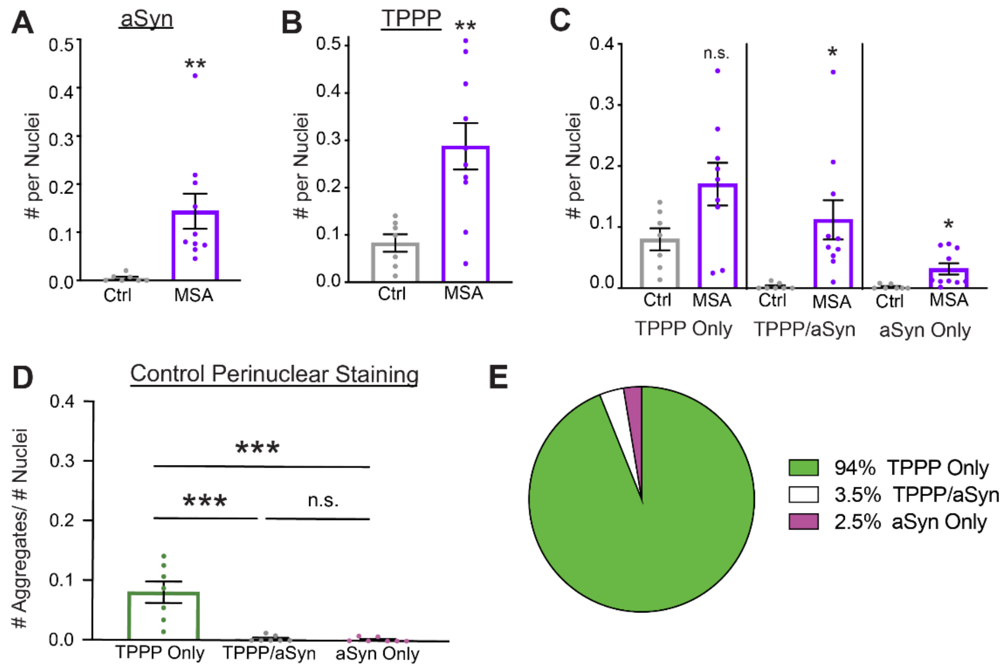


**Fig. S4. aSyn and TPPP perinuclear staining in MSA.**
(A) Density of aSyn perinuclear staining, normalized by the number of DAPI-positive nuclei. N = 7, 10 patients; 2 images per subject. Student's *t*-test: **P < 0.01.
(B) Density of all types of TPPP perinuclear staining for the aSyn images, normalized by the number of DAPI-positive nuclei. n = 7, 10 patients; 2 images per subject. Student's *t*-test: **P < 0.01.
(C) Comparison between control and MSA SN for each type of staining: TPPP only, aSyn only, and TPPP/aSyn colocalized. Normalized by DAPI-positive nuclei. Student's *t*-test: *P < 0.05, n.s. P = 0.054.
(D) In control brains, comparison of the three types of TPPP/aSyn colocalizations. Normalized by DAPI-positive nuclei. n = 7 control; 2 images per subject. One-way ANOVA: *P < 0.05, **P < 0.01, ***P < 0.001, n.s. P = 0.999.
(E) Pie chart illustrating percentages/distribution of all TPPP/aSyn perinuclear staining in control SN.

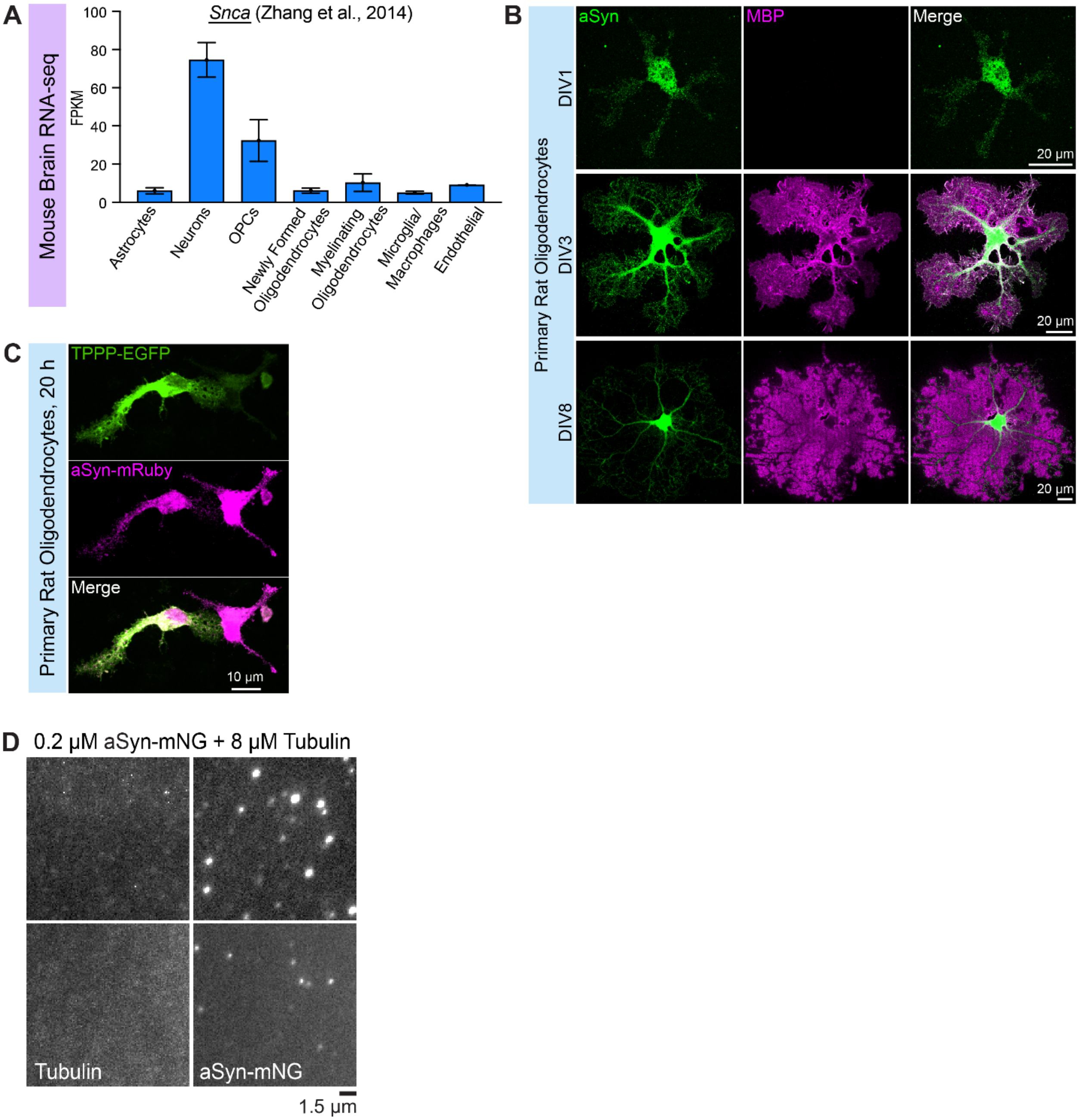


**Fig. S5. aSyn expression in oligodendrocytes and liquid condensate behavior.**
(A) *Snca* (aSyn) mRNA expression in a bulk RNA-seq database of immunopanned brain cells from *Mus musculus* brains (*10*). FPKM (fragments per kilobase of transcript per million mapped reads).
(B) Primary rat oligodendrocyte immunostaining of endogenous aSyn and MBP (myelin basic protein) at different developmental time points or days *in vitro* (DIV1, DIV3, DIV8).
(C) Primary rat oligodendrocytes expressing TPPP-EGFP and aSyn-mRuby. Oligodendrocytes were electroporated with expression constructs, then differentiated for 20 h.
(D) TIRF microscopy images of tubulin and aSyn without TPPP. 0.2 μM aSyn-mNG, 8 μM tubulin-CF640R in BRB80.

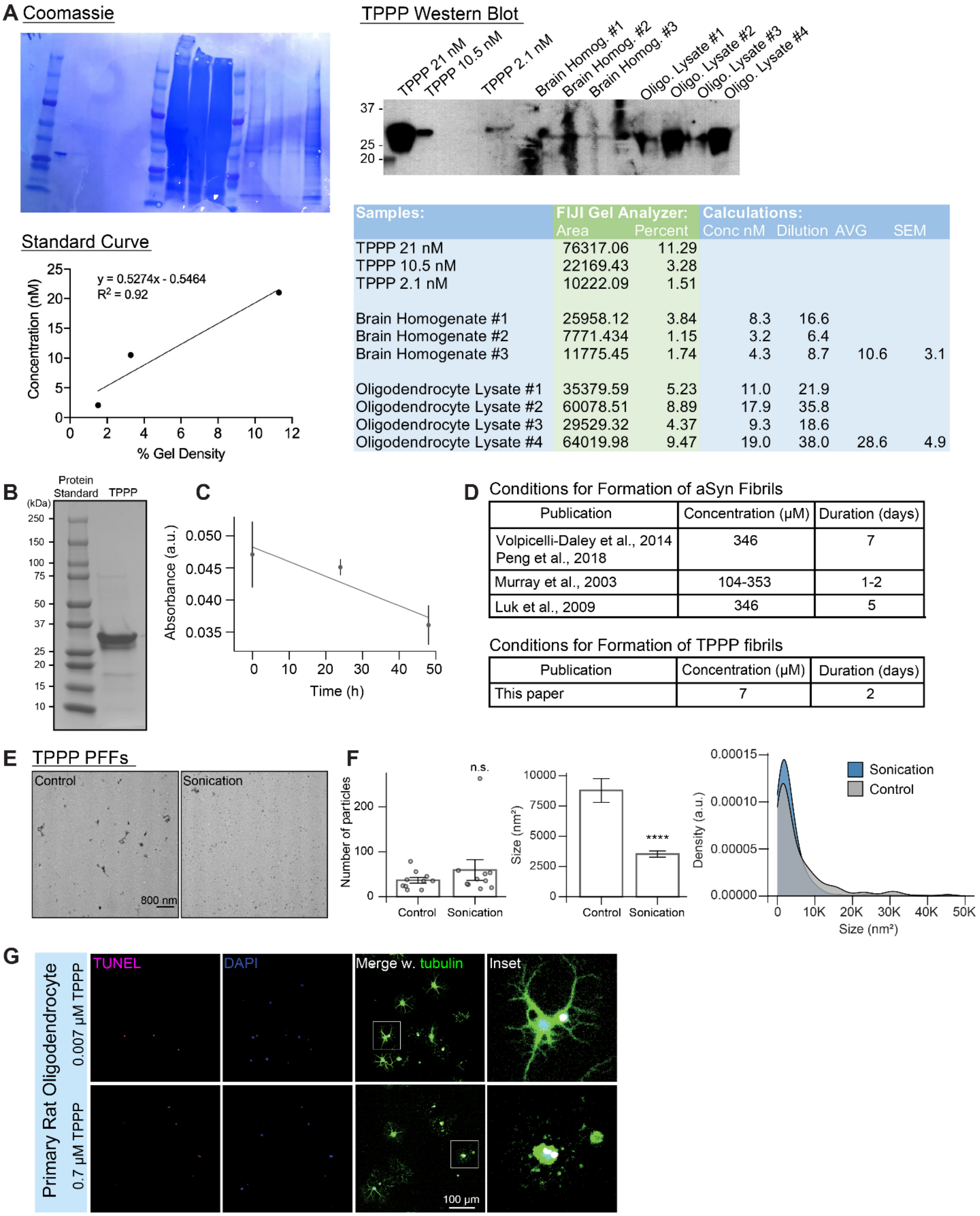


**Fig. S6. TPPP endogenous level calculations and generation of TPPP PFFs.**
(A) Quantification of endogenous TPPP concentrations in rodent brain homogenates and primary oligodendrocyte lysates. A standard curve was generated from the Western

blot densities of unlabeled recombinant TPPP and used to calculate TPPP concentrations.
(B) Purification of recombinant unlabeled TPPP used to generate TPPP PFFs.
(C) Absorbance of TPPP PFF reactions after different timepoints of incubation at 37°C and shaking. n = 3 experiments.
(D) Comparison of published concentrations and durations of shaking used to generate aSyn PFFs.
(E) Representative TEM micrographs of TPPP PFFs before and after sonication.
(F) The mean size of TPPP PFFs decreased ~2.5-fold from 8782 $nm^2$ to 3526 $nm^2$ after sonication. Assuming a circular shape, sonicated PFFs have diameter ~67 nm. n = 365 control, 592 sonicated PFFs. Kolmogorov-Smirnov test; ****P < 0.0001.
(G) TUNEL staining of differentiated primary rat oligodendrocytes treated with 0.007 µM and 0.7 µM TPPP PFFs. Black arrowheads indicate TUNEL-positive cells. White triangles indicate DAPI-negative TUNEL-negative dead “shadow” cells in which the nuclei have detached.
All error bars represent mean ± SEM.

| Subject ID | PMI (hours) | Age | Race | Sex | Sequential Extraction (Y/N) |
|---|---|---|---|---|---|
| **MSA** | | | | | |
| S03266[a] | 23.95 | 65 | White | Male | N |
| S05842[a] | 16.82 | 70 | White | Female | N |
| S09164 | 14.75 | 74 | White | Female | N |
| S11347 | 9.92 | 75 | White | Male | Y |
| S14396 | 18.3 | 76 | White | Male | N |
| S15282 | 13 | 65 | White | Female | N |
| S15799 | 28.25 | 69 | White | Female | N |
| S16023 | 6.5 | 57 | White | Female | Y |
| S16219 | 16.28 | 61 | White | Female | Y |
| S19671 | 18.58 | 80 | White | Female | N |
| **Control** | | | | | |
| S01733 | 16.8 | 61 | White | Female | Y |
| S02276 | 16.62 | 53 | White | Female | N |
| S06589 | 12.53 | 60 | White | Female | Y |
| S08610 | 23.38 | 78 | White | Female | Y |
| S08682 | 22.5 | 66 | White | Male | N |
| S12016 | 30.18 | 65 | White | Female | N |
| S16830 | 23.92 | 68 | White | Female | N |

[a]These cases do not have TPPP accumulations similar to the other 8 cases.

**Table S1.** Patient brain sample IDs and data.

**Movie S1.** TPPP liquid condensates nucleate microtubules.

**Movie S2.** FRAP of oligodendrocytes expressing punctate TPPP-EGFP.

**Movie S3.** FRAP of oligodendrocytes expressing aggregate filamentous TPPP-EGFP.

**Movie S4.** Control and TPPP PFF-treated oligodendrocytes imaged with annexin V dye.
